# Chromosomal rearrangements and instability caused by LINE-1 retrotransposition

**DOI:** 10.1101/2024.12.14.628481

**Authors:** Carlos Mendez-Dorantes, Xi Zeng, Jennifer A. Karlow, Phillip Schofield, Serafina Turner, Matthew Leventhal, Jupiter Kalinowski, Jinyu Wang, Sonia Zumalave, Jose M. C. Tubio, Eunjung Alice Lee, Kathleen H. Burns, Cheng-Zhong Zhang

## Abstract

LINE-1 (L1) retrotransposition is widespread in many cancers, especially those with a high burden of chromosomal rearrangements. However, whether and to what degree L1 activity directly impacts genome integrity is unclear. Here, we apply whole-genome sequencing to experimental models of L1 expression to comprehensively define the spectrum of genomic changes caused by L1. Combining the analyses of experimental models of L1 induction and of cancer genomes, we demonstrate that L1 retrotransposition can directly generate reciprocal translocations, genomic DNA inversions, and foldback rearrangements resulting from illegitimate recombination of double-strand DNA ends generated by L1-encoded ORF2p. We further show that L1-induced rearrangements can produce unstable chromosomes that fuel the acquisition of complex rearrangements, large segmental copy-number alterations, and genetic heterogeneity through breakage-fusion bridge cycles or DNA fragmentation. Together, these findings suggest L1 as a potent mutagenic force capable of driving genome evolution in cancers.

## INTRODUCTION

Long INterspersed Element 1 (LINE-1 or L1) is the only autonomous retrotransposon in the human genome^1–3^. L1 encodes two proteins that are required for retrotransposition^4,5^: the open reading frame 1 protein (ORF1p), an RNA-binding protein^6,7^; and the open reading frame 2 protein (ORF2p) with both endonuclease (EN)^8^ and reverse transcriptase (RT) activities^9,10^. Retrotransposition occurs via target-primed reverse transcription (TPRT)^11^ (**Figure 1A**): ORF2p first nicks genomic DNA (3′-AA/TTTT-5′), initiating a primer-template duplex between the T-rich sequence and the poly-A tail of the mRNA; ORF2p then reverse transcribes the mRNA to create a cDNA. These steps have been recapitulated *in vitro*^10,12,13^. The cDNA is eventually converted to double-stranded DNA (dsDNA), but the mechanism of this conversion is largely unknown^9,14^.

**Figure 1.**
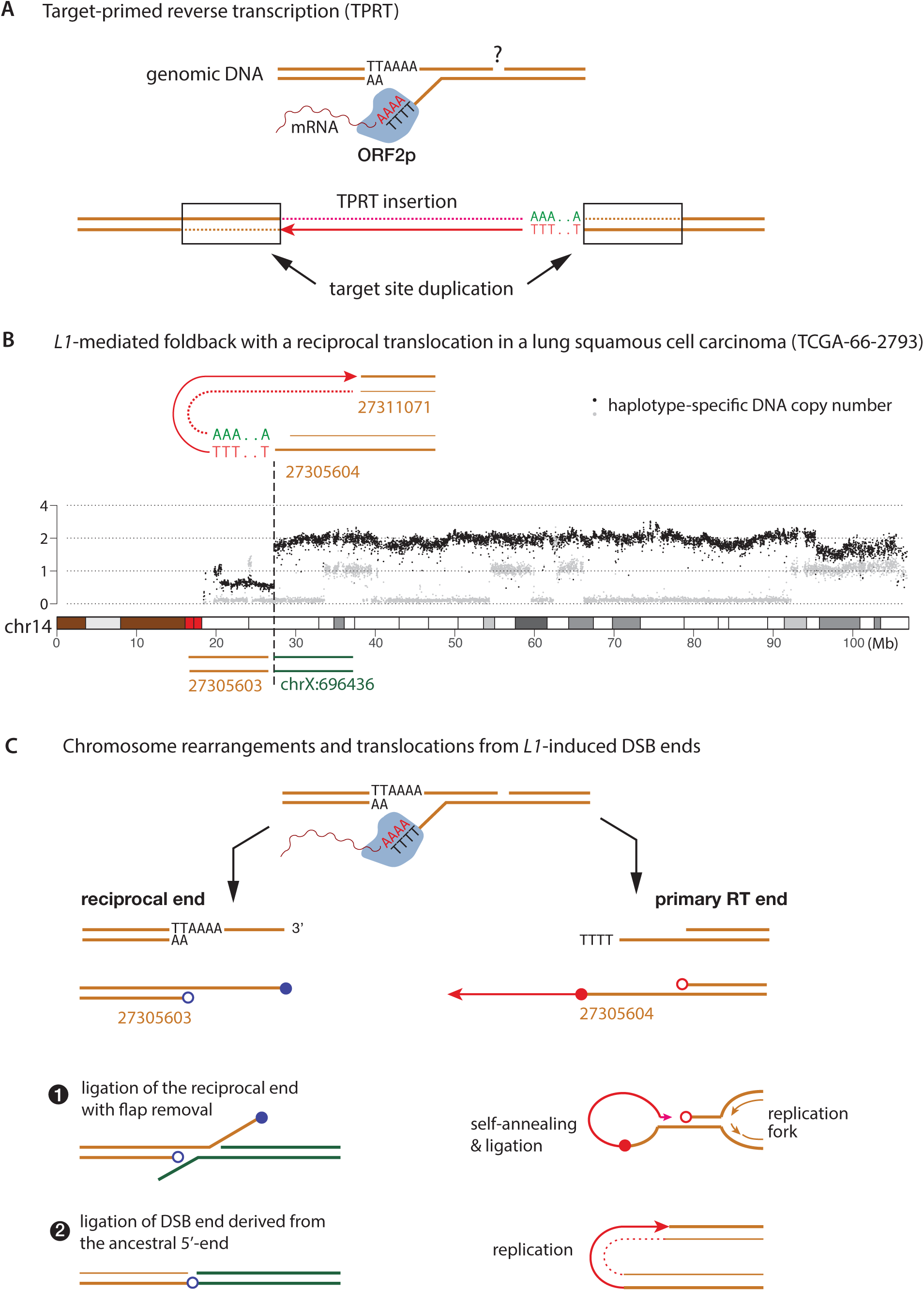
An example of L1-mediated rearrangement indicating DNA double-strand break. **A.** Target-primed reverse transcription (TPRT) mediated by L1 ORF2p. (Top) A duplex between the genomic DNA (brown solid lines) and the mRNA template (wiggly line) formed after ORF2p nicks the bottom DNA strand at an A/T rich target site (TT|AAAA). A possible nick of the top strand is marked with a question mark. (Bottom) The final insertion outcome after reverse transcription (RT) of the mRNA sequence. The first cDNA strand generated by RT is shown as a solid red line with arrowhead pointing to the 3’-end; the dotted line represents the complementary strand (2^nd^ strand). Duplicated target sequences are outlined. **B.** L1-mediated DNA rearrangement identified in a lung squamous cell carcinoma. Black and gray dots represent haplotype-specific DNA copy number. A truncated L1 integration is identified at a foldback junction between two breakpoints near 27.3Mb (27,305,604 and 27,311,071). There is a translocation between a 3^rd^ breakpoint on the reciprocal side (27,305,603) and chrX. The breakpoints are co-localized with the copy-number transition of the homolog whose copy number is shown in black. In the original paper where this example was presented, the left translocation breakpoint was not reported; and the second foldback breakpoint (27,311,071) was suggested to derive from an independent DNA break end. We suggest that all three breakpoints derive from ancestral ssDNA ends generated during a single TPRT event. **C.** A proposed model for generating the foldback and the translocation shown in **B** in a single TPRT event. Nicking of both DNA strands creates two double-strand break (DSB) ends: The right DSB end with a 3’-end extended by ORF2p RT is hereafter referred to as the primary RT (DSB) end; filled and open red circles represent the 3’- and 5’-ends of the primary RT end. The left DSB end is referred to as the reciprocal end; filled and open blue circles represent the 3’-and 5’-ends of the reciprocal DSB end. If the primary RT and the reciprocal ends are joined together, it will create an insertion outcome as shown in **A**. For the example shown in **B**, the L1 insertion sequence identifies the breakpoint at 27,305,604 to be the ancestral 3’- primary RT end. Based on proximity, we infer the translocation breakpoint 27,305,603 to have derived from the reciprocal 5’-end. The foldback junction on the right arises by replication/fusion of the primary RT end after TPRT; the translocation junction on the left arises from ligation of the reciprocal end to a distal DSB end on chrX. In this study, we use the sequence features of TPRT insertions (poly-A and ORF2p EN target sequence) to identify breakpoints derived from the ancestral 3’-primary RT ends and infer the relationship between the other adjacent breakpoints and ancestral DNA ends (the 5’-primary RT end and the 3’- or 5’- reciprocal ends) based on breakpoint proximity.

L1 elements are epigenetically silenced during early development but can be activated in somatic cells^15^ and in human malignancies^16,17^. Somatic L1 activation has been demonstrated by the hypomethylation of L1 promoters, the presence of ORF1p in malignant tissues^18–20^, and somatic L1 insertions in both pre-cancerous lesions^21^ and advanced cancers^22–27^. Intriguingly, analyses of cancer genomes have revealed not only L1 insertions, but also L1 sequences at long-range rearrangement junctions^27^. Although it has been shown that L1 expression causes DNA breakage^28^ and elicits DNA damage response^29^, the genomic observations of L1-mediated rearrangements raise three questions about L1 retrotransposition and genome integrity. How does aberrant or incomplete L1 integration contribute to chromosomal rearrangements? What kinds of rearrangements can arise from L1 retrotransposition? And what is the impact of L1 activity on genome evolution beyond canonical insertions?

To address these questions, we have developed experimental systems with inducible L1 expression to analyze the full spectrum of genomic alterations of L1 activation. Using a combination of short- and long-read whole genome sequencing, we identify three classes of L1-induced chromosome rearrangements: reciprocal translocations, small inversions or inverted duplications, and foldback rearrangements. These outcomes recapitulate observations in cancer genomes^27,30^. Mechanistically, we found all these rearrangement outcomes can be explained by recombination, processing, or replication of DNA double-strand break ends generated during TPRT, highlighting a dynamic interplay between retrotransposition, DNA damage, and genome plasticity. We further show how L1-induced rearrangements produce unstable chromosomes that give rise to large segmental copy-number alterations and instigate further chromosomal instability. Together, these findings substantially expand the repertoire of L1-induced genomic alterations and suggest a profound role of L1 activation in the evolution of cancer genome complexity and heterogeneity.

## RESULTS

### Genome evidence of L1-induced DNA double-strand breakage

Although the nicking of genomic DNA by ORF2p EN prior to initiate TPRT is well-established (**Figure 1A**), how and when the second strand is nicked for the resolution of a new insertion is poorly characterized^9,10,12,13^; therefore, it is unclear whether retrotransposition directly generates DSB ends. We report genomic evidence of a L1-induced DSB that was provided by an observation of L1-mediated rearrangement in a lung cancer reported previously^27^ (**Figure 1B**).

In this example, a truncated, polyadenylated L1 sequence was identified at a foldback junction near 27.3Mb on chr14 (shown above the copy-number plot); we further identified a translocation breakpoint (chr14:27,305,603, shown below the copy-number plot) that was not reported in the original study. These three breakpoints coincide with a copy-number transition on the homolog whose copy number was shown in black. Based on the L1 insertion and the proximity between these breakpoints, we infer that all the breakpoints originate from an ancestral DNA double-strand break (DSB) generated during TPRT (**Figure 1C**). The integrated L1 sequence identifies the breakpoint at 27,305,604 to be the ancestral 3’-end that was extended by ORF2p RT (i.e., the primary RT end); based on proximity, we infer the translocation breakpoint at 27,305,603 to have derived from the reciprocal end (likely the 5’-end). We further infer the second foldback breakpoint (27,311,071) to have derived from the resected 5’-end of the primary RT end; notably, the generation of a foldback junction indicates a double-stranded primary RT end in the ancestral DNA^31^. Thus, the two rearrangements indicate different recombination outcomes of two ancestral DSB ends generated during ORF2p-mediated TPRT. Therefore, TPRT can generate DSB and L1-mediated rearrangements may be initiated by L1-induced DSB.

### Experimental evidence of L1-induced DNA double-strand breakage

We next sought direct experimental evidence that TPRT can cause DSB. Prior studies indicated that p53 inactivation is required for cell proliferation after L1 expression^29^. We therefore established a Tet-On system of inducible L1 expression in p53-/-, hTERT immortalized RPE-1 cells (**Figure 2A, Methods**). To distinguish insertions or rearrangements involving the experimentally induced L1 from those involving endogenous L1 sequences, we used a codon-optimized sequence of human L1 (L1 ORFeus)^32^.

**Figure 2.**
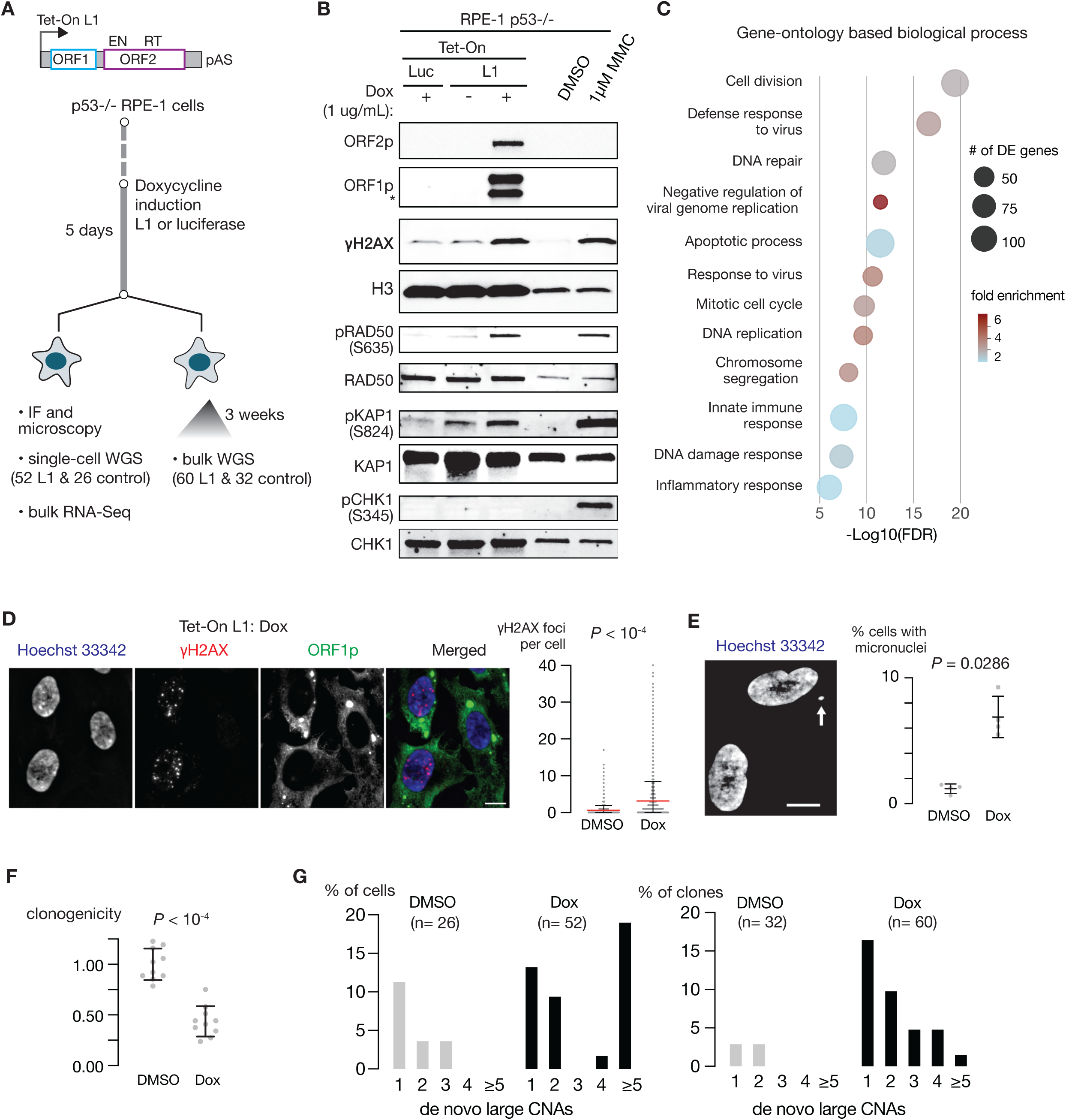
L1 expression causes extensive DNA damage including double-strand breaks. **A.** Experimental and analytical workflow. Expression of a codon-optimized L1 (ORFeus) is induced in p53-null RPE-1 cells for five days using a Tet-On expression system. Cells with L1 or luciferase expression induced with Doxycycline, or treated with DMSO (no induction) are used for subsequent analyses. **B.** Immunoblots of L1 encoded proteins ORF1p and ORF2p and markers of DNA damage including γ-H2Ax, pRAD50 (S635), pKAP1 (S824), pCHK1 (S345) from whole cell lysates after L1 induction and control experiments: Luc for induced luciferase expression and DMSO for no induction; both are negative controls. Cells treated with 1 µM mitomycin C (MMC) serve as a positive control. See also **Figure S1A** and **S1B**. **C.** Biological processes inferred from Gene Ontology analysis of differential gene expression due to L1 expression. Differentially expressed (DE) genes are identified by a comparison of p53-/- RPE-1 cells with Dox-induced L1 expression and with Dox-induced luciferase expression (*n* = 3 technical replicates for each condition), using a threshold of adjusted *P* < 0.05. The size of each circle reflects the number of DE genes associated with each process; the color shade reflects the enrichment of DE genes (fold change of gene count) in each process. See also **Figure S1C**. **D.** Increased γ-H2Ax foci after L1 induction. Left: representative images of γ-H2AX foci in cells after L1 expression (Bar scale: 10µm). Right: Quantification of γ-H2AX foci per cell with (*N* = 2,889 cells) and without (*N* = 2,274 cells) L1 induction (*n* = 2 experiments). *P* < 0.0001; two-tailed Mann-Whitney U-test. See **Figure S1D** for data on 53BP1 foci. **E.** Increased micronucleation after L1 induction. Left: an example of a micronucleus in a cell after L1 induction (Bar scale:10µm). Right: Quantification of cells with micronuclei after L1 induction (*n* = 4 independent experiments; DMSO: 1,820 cells; Dox: 1,223 cells; *P* = 0.0286; two-tailed Mann-Whitney U-test). **F.** Reduced clonogenicity of cells after L1 induction (5-day treatment of Doxycycline) in comparison to control cells (5-day treatment of DMSO). *P* <0.0001; two-tailed Mann-Whitney U-test. See also **Figure S1A**. **G.** Large segmental copy-number alterations in single cells (left) and progeny clones derived from single cells (right) after L1 induction. Shown are the percentage of single cells or single-cell derived clones that harbor 1, 2, 3, 4 or ≥ 5 de novo DNA copy-number alterations ≥ 5Mb assessed from 0.1× whole-genome sequencing data. *P* = 0.0211 for single cells, *P* = 0.0011 for single-cell derived clones; Fisher’s exact test. See also **Figure S2A**.

We first confirmed the specific expression of L1 induced by Doxycycline (Dox) (**Figure 2B**, **Figure S1A**). In cancer genomes, L1-mediated rearrangements were often associated with a high burden (10^2^) of L1 insertions^30^, which can reflect either a long duration of L1 expression or an episodic burst of high L1 expression. For our experimental L1 induction, we chose to induce potent L1 expression for a short period (5 days) to increase the chance of recapitulating the processes that lead to L1-mediated rearrangements. After L1 induction, we observed elevation of molecular markers of DSB, including γH2AX and other targets of the ATM kinase (pRAD50 and pKAP1) (**Figure 2B**, **Figure S1B**). Elevated DNA damage was also indicated by the upregulation of genes associated with DNA damage response and DNA repair (**Figure 2C**, **Figure S1C**, **Table S1, S2**).

At the cellular level, DSB generation after L1 induction was demonstrated by the increase of γH2AX (**Figure 2D)** and 53BP1 nuclear foci (**Figure S1D-E**) indicating nascent DSBs, and the formation of micronuclei (**Figure 2E**) from acentric chromosome fragments due to unrepaired DSBs. Using Tet-On inducible L1 mutants lacking endonuclease (EN) or reverse transcriptase (RT) activities established in U2OS cells, we confirmed that both EN and RT activities contribute to DSB formation, with ORF2p EN being the dominant contributor (**Figure S1F-G**).

Finally, L1 induced DSB was demonstrated by the presence of large segmental copy-number alterations (CNAs) in both single cells harvested immediately after L1 induction and their progeny clones (**Methods**). In comparison to control cells or clones, L1 induction resulted in a 50% reduction of clonogenicity (**Figure 2F**) and a higher burden of large segmental (5Mb or above) CNAs (**Figure 2G**, **Figure S2A**). (The observation of more single cells with a high CNA burden (>4 large CNAs) than single-cell derived clones likely reflects reduced clonogenicity of cells with a higher CNA burden or a larger number of unrepaired DSBs.) The elevated burden of large CNAs after L1 expression was also recapitulated in another L1 expression system using an L1-GFP reporter^33^ (**Methods, Figure S2B-D**).

Together, these results demonstrate that L1 expression causes DNA DSBs and leads to the acquisition and accumulation of large segmental CNAs.

### Identification of L1-induced genomic alterations by whole-genome sequencing

We next sought to determine the full spectrum of L1-induced genomic alterations by whole-genome sequencing (WGS) (**Methods**). From the Dox-induced L1 expression system, we generated short-read WGS data on 28 single cells after 5 days of Dox-induced L1 expression (Dox L1 single cells), 31 clones expanded from single cells after L1 expression (Dox L1 clones), 12 single cells with DMSO treatment, and 10 control clones expanded from single cells with DMSO treatment. Using the L1 GFP reporter, we generated short-read WGS data on 29 clones derived from GFP+ single cells (GFP L1 clones) and 5 clones derived from GFP-single cells. To resolve the complete sequences of both insertion and rearrangement junctions, we also generated PacBio HiFi sequencing (median 15x) on 38 L1 clones and one control clone.

We performed comprehensive mutation discovery (single-nucleotide substitutions, short insertions/deletions, chromosomal rearrangements, and DNA copy-number alterations) from short-read data using bioinformatic workflows described previously^34^. L1 expression can lead to both L1 insertions and insertions of non-autonomous transposons (*Alu* and *SVA*) or endogenous mRNAs (processed pseudogene insertions)^35,36^. Insertions of L1 or other retrotransposons were detected using xTea^37^; insertions of processed pseudogenes were detected using sideRETRO^38^. All insertions were also detected independently as part of DNA rearrangement discovery and the complete insertion sequences were resolved by long reads when available.

Here, we focus on insertions associated with or accompanied by genomic DNA rearrangements (**Table S3-S6**); the complete analysis of L1-induced insertional mutagenesis is presented in a separate paper. We note that many pseudogene insertions were detected in Dox L1 cells and clones. This may be partially due to either the high level of L1 induction in our experimental system or the usage of a codon-optimized L1 that may have reduced affinity with the *cis* L1 mRNA. However, for the identification and mechanistic interpretation of chromosomal rearrangements from DSB ends generated during TPRT, the inserted sequences are of less importance than the general features of the TPRT insertions, i.e., polyadenylation, truncation, or inversion. We therefore refer to both L1 and processed cDNA insertions as TPRT insertions.

To complement the genomic analysis of experimentally induced L1 expression, we further analyzed L1-mediated rearrangement junctions in 10 cancer genomes for which both short and long-read sequencing data were available^30^. For the cancer genomes, we further calculated haplotype-specific copy number^39^ and annotated copy-number changes at L1-mediated rearrangement junctions that were reported previously^30^ (**Table S7**). The haplotype-specific DNA copy number provided phasing information that can relate L1-mediated rearrangement breakpoints to segmental copy-number alterations on the same homologous chromosome.

In the following sections, we present observations from both cancer genomes and experimentally derived cells or clones that represent three classes of rearrangements of the genomic DNA sequences near the integration sites of TPRT insertions.

### Recombination between L1-induced DSBs creates reciprocal translocations

The example shown in **Figure 1B** suggests that a single TPRT event can generate two DSB ends that undergo different recombination events. The simplest outcome of such events is an exchange between two pairs of DSB ends that leads to two reciprocal translocations^40,41^. Such events can create either two stable derivative chromosomes (**Figure 3A**) or two unstable chromosomes (one dicentric, one acentric); the latter outcome will be considered later.

**Figure 3.**
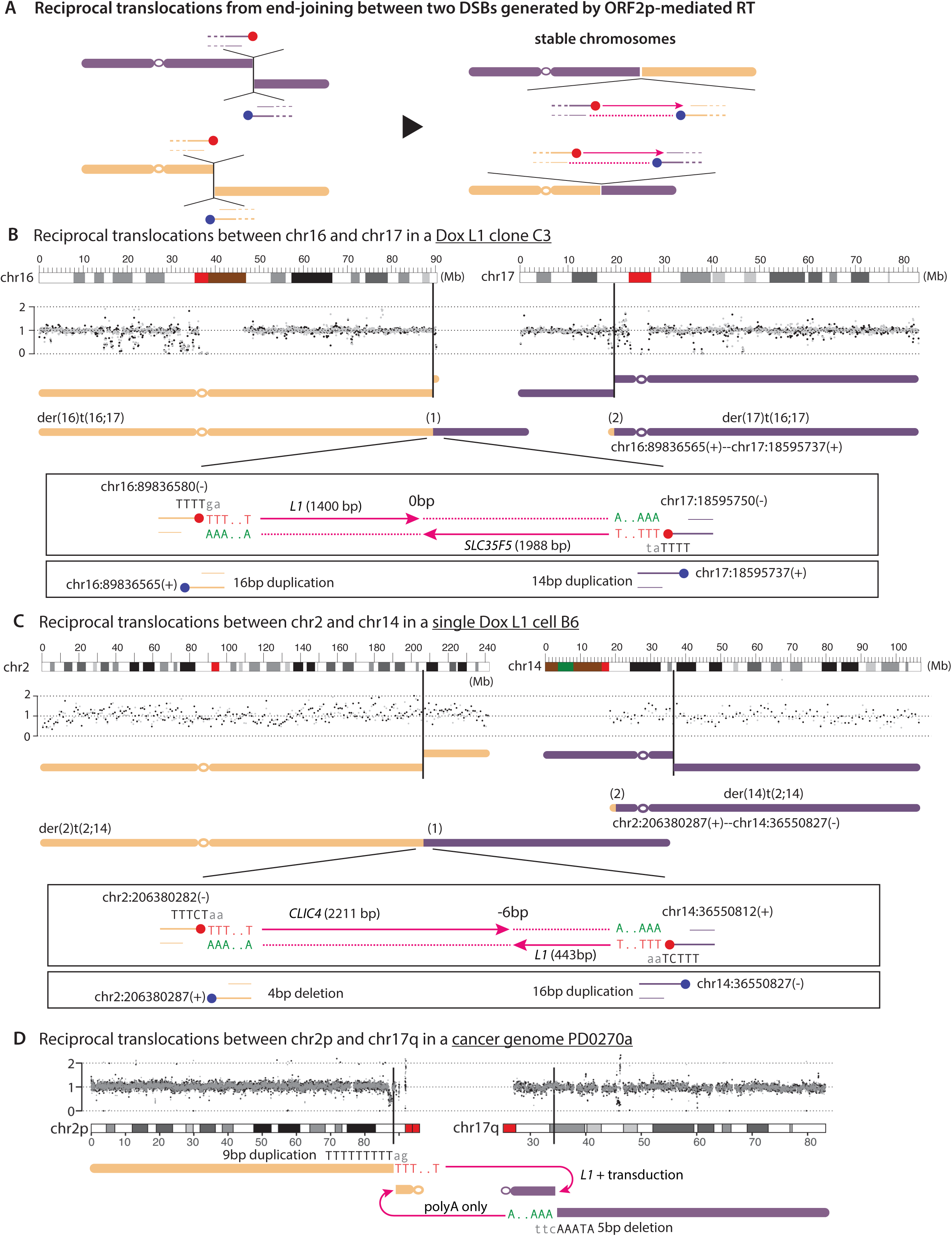
Reciprocal translocations from recombination between ORF2p-induced DSBs. **A.** Schematic diagram of reciprocal translocations from an exchange between two pairs of DSB ends generated by ORF2p: 3’ primary RT ends are marked with filled red circles; the 3’ reciprocal ends are marked with filled blue circles. TPRT insertions (red lines with arrowheads) may be retained in separate translocation junctions (shown here) or in a single junction (model not shown; examples shown in **B** and **C**). The translocations can create two stable chromosomes (shown here) or two unstable chromosomes (one acentric and one dicentric) that will be discussed later (**Figure 7B**). **B.** Reciprocal translocations between chr16 and chr17 in a clone (C3) derived from a single RPE-1 cell after Dox-induced L1 expression. Gray and black dots represent normalized DNA copy number (90kb bins) of two parental chromosomes, showing no allelic imbalance as expected for reciprocal translocations. The breakpoints at chr16:89,836,580(-) and chr17:18,595,750(-), both located within potential ORF2p EN target sequences (5’-TTTT|ga-3’ on chr16, 5’-TTTT|ata-3’ on chr17) and adjacent to the 3’ end (poly-A/T) of TPRT insertions, derive from the primary RT ends (hereafter “primary RT breakpoints”); they are joined together to form the translocated chromosome der(16)t(16;17). The other translocated chromosome der(17)t(16;17) is formed by recombination between chr16:89,836,565(+) and chr17:18,595,737(+); we infer that these breakpoints derive from the reciprocal ends (reciprocal breakpoints). There are partial duplications between the primary RT and the reciprocal breakpoints at both loci (16bp at the site from chr16, 14bp at the site from chr17) that recapitulate the “target-site duplication” feature of TPRT insertions. There is no microhomology or insertion at the junction between the two RT breakpoints (‘0bp’), and a 3bp microhomology between the reciprocal breakpoints. Note: We use ‘-’ to denote breakpoints corresponding to the right boundary of a rearranged sequence when aligned to the reference, ‘+’ for breakpoints corresponding to the left boundary of a rearranged sequence when aligned to the reference. The breakpoint coordinates denote the 1-based positions of the last retained nucleotides on both ends. We use black, uppercase letters to denote nucleotides that are retained in the rearrangement junction and gray, lowercase letters for the adjacent nucleotides in the reference. The number of base pairs at translocation junctions denotes insertions (plus), blunt joining (0bp), or homology (minus). **C.** Reciprocal translocations between chr2 and chr14 in a single cell (B6) after Dox-induced L1 expression. Junctions and breakpoints are annotated similarly as in **B**. The chr2 breakpoints show a 4bp deletion and the chr14 breakpoints show a 16bp duplication. There is a 6bp homology between the breakpoints derived from the RT ends (‘-6bp’), and no homology or insertion between the breakpoints derived from the reciprocal ends. **D.** Reciprocal translocations between chr2 and chr17 in a cancer genome (PD0270a). The chr2 breakpoints display a 9bp duplication and the chr17 breakpoints display a 5bp deletion.

From samples with experimentally induced L1 expression, we identified two examples that recapitulated the stable reciprocal translocation outcome. In the first example from a Dox L1 clone (**Figure 3B**), reciprocal translocations between chr16 and chr17 created two derivative chromosomes with no DNA copy-number change of either homolog. The origin of all four rearrangement breakpoints from DSB ends generated by L1 ORF2p is established by three sequence features of the breakpoints: (1) polyadenylated insertions that identify the ancestral 3′ primary RT end; (2) ORF2p EN target sequences at the primary RT breakpoint; and (3) small duplications shared between the primary RT and the reciprocal breakpoints (16bps on chr16 and 14bps on chr17) that resemble the target site duplication (TSD) feature of canonical TPRT insertions. A similar translocation outcome involving chr2 and chr14 was also observed in a Dox L1 single cell (**Figure 3C**).

From 10 cancer genomes, we identified 12 instances of reciprocal translocations with polyadenylated TPRT insertions (**Table S7,** ‘2x balanced’ in Column I). Eight of these contained two polyadenylated insertions indicating recombination between two pairs of L1-induced DSB ends as shown in **Figure 3A**. In two instances, both insertions were retained in one junction; in six instances (one of which is shown in **Figure 3D**), each junction retained one insertion. In the remaining four instances, polyadenylated TPRT insertions were identified at one break site but not the other. These translocations may reflect recombination between L1-induced and L1-independent DSB ends, which was recently demonstrated experimentally^42^. We further identified two large para-centric inversions (PD0277a, chr5:30.5-32.2Mb; PD0307a, chr5:19.0-20.5Mb; **Figure S3A**) that arose from an exchange between two pairs of L1-induced DSB ends on a single chromosome arm. Thus, the observations from both experimental L1 clones and cancer genomes demonstrate the outcomes of recombination between DSB ends from two target sites.

We further identified multi-way balanced translocations known as chromoplexy. In an example from a cancer genome (**Figure S3B**), we identified four translocations involving three pairs of reciprocal breakpoints and two single breakpoints from five chromosomes. Polyadenylated TPRT insertions identified two pairs of breakpoints that were generated by L1-induced DSB. In an example from a Dox L1 clone (**Figure S3C**), we identified three translocations between two pairs of reciprocal breakpoints and two single breakpoints from four chromosomes. In the latter example, the two breakpoints on chr18 each contained a cDNA fragment of the *OSBPL8* mRNA that recapitulate the intermediate state predicted by the twin-priming model^3,43^. Had these two breakpoints be ligated together, it would have resulted in a 5’-inverted pseudogene copy of *OSBPL8*. Given that only the 3’-end can be extended by reverse transcription, the observation of inverted *OSBPL8* cDNA insertions at two separate translocation junctions indicates that both the primary RT end and its reciprocal end were DSB ends at the time of translocation.

In summary, we observed three rearrangement outcomes from illegitimate recombination between L1-induced DSB ends: two-way reciprocal translocations, intra-chromosomal inversions, and chromoplexy.

### Local sequence rearrangements suggest remodeling of L1-induced DSB ends

The second class of L1-induced rearrangements consists of inversions or inverted duplications of genomic DNA sequences near the site of L1 integration (**Figure 4**). Such rearrangements can be explained by self-annealing of ssDNA flaps after DSB end resection^44^ (**Figure 4A**). Cleavage of the flap or hairpin can result in either DNA deletion (1) or DNA inversion (2). If the self-annealed 3’-end primes DNA synthesis, it can also create an inverted duplication (3), as suggested in a recent study^45^. Small genomic deletions at the site of L1 integration have been observed in both experimental systems^46,47^ and somatic human cells^48^. Consistent with the model of DSB end remodeling, most L1-mediated deletions in cancer genomes are shorter than 10kb^27^ (also see **Figure S5A**) and within the range of DSB end resection^49,50^. The model of DSB end remodeling is further supported by observations of both the inversion and the inverted duplication outcomes in our experimental system and in cancer genomes (**Figure 4B-E**).

**Figure 4.**
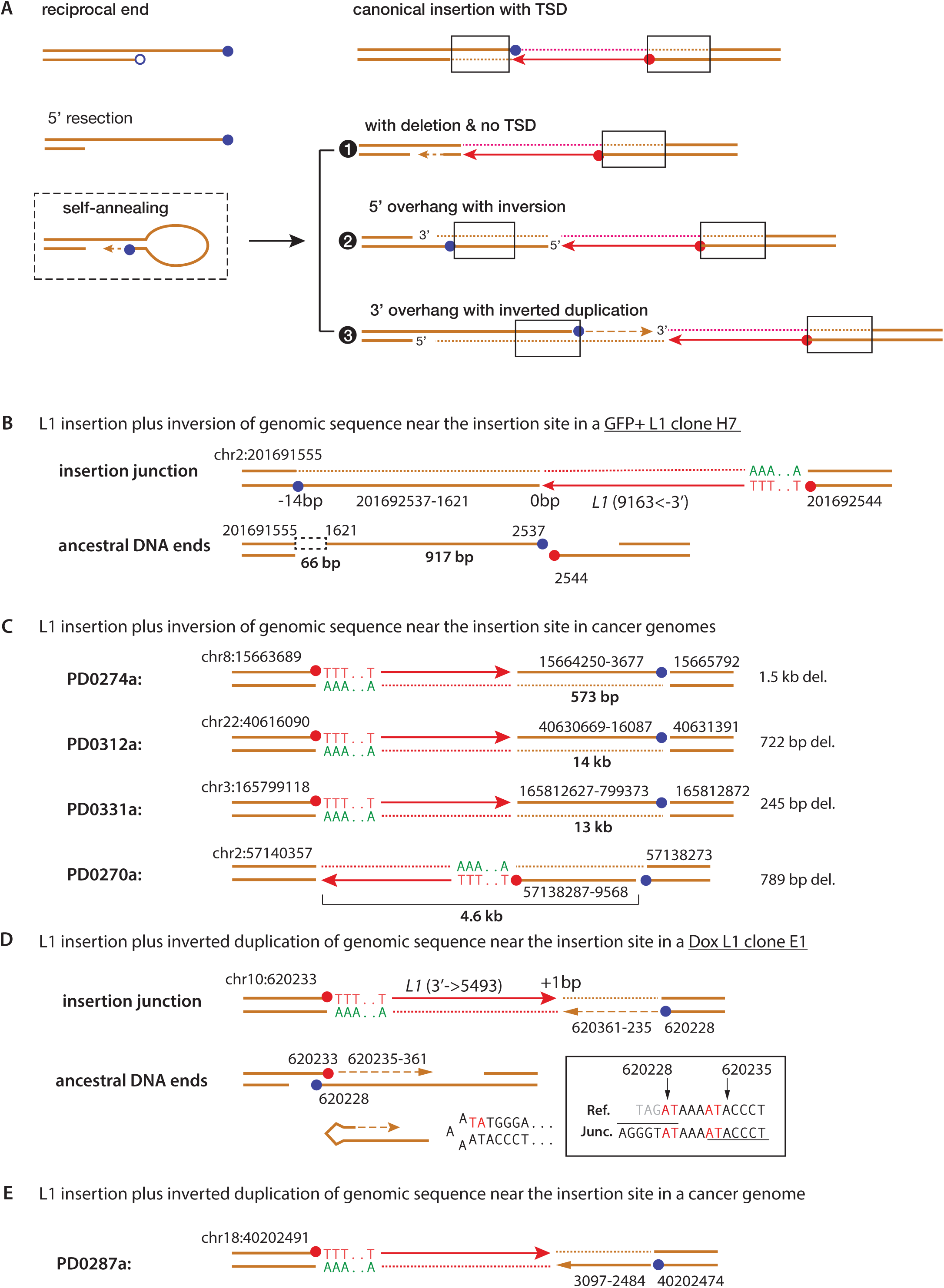
Inversion or inverted duplication of genomic DNA at ORF2p-induced DSB ends. **A.** A proposed model of local genomic DNA rearrangements due to remodeling of ORF2p-induced DSB ends. Left: resection of the 5’-end (open circle) of the reciprocal DSB end creates a large 3’-overhang, which can anneal with itself using internal microhomologies. Right: ligation between the remodeled reciprocal end and the primary RT end can create three possible outcomes: (1) deletion; (2) inversion; (3) inverted duplication. Duplication between the primary RT end (red circle) and the reciprocal end (blue circle) is highlighted with boxes. **B.** An example of L1 insertion with genomic DNA inversion on the reciprocal end identified in a GFP+ L1 clone (H7). Note the inversion of the reciprocal 3’-end (blue filled circle) from the top strand to the bottom strand. **C.** Three instances of L1 insertion with genomic DNA inversion on the reciprocal end and one instance of L1 insertion accompanied by inversion on the primary RT end identified in four different cancer genomes. Note the overlap between the primary RT and reciprocal breakpoints (red and blue circles) that resemble TSD at TPRT target sites in the examples from PD0274a (13bp), PD0312a (4bp), and PD0270a (15bp). The inversion in PD0331a shows a 254bp deletion at the target DSB site. The sizes of the inverted sequences are annotated at the bottom; for the example from PD0270, this includes both the genomic DNA and the L1 insertion (3,275bp). The sizes of deletions represent the gap between the inverted sequence and the proximal breakpoint. **D.** An example of L1 insertion with an inverted duplication of the genomic sequence on the reciprocal end identified in a Dox L1 clone (E1). The sequence at the inversion junction (highlighted in the box) suggests 2bp homology (TA/AT) that primes DNA synthesis. **E.** An example of L1 insertion with an inverted duplication of the genomic sequence on the reciprocal end identified in a cancer genome (PD0287a). There is a 2bp homology between the inverted breakpoints. Note the duplication feature between the primary RT (red circles) and the reciprocal breakpoints (blue circles) in both **D** and **E**.

**Figure 4B** shows an inversion outcome identified in an experimental L1 clone. Here, the insertion junction contained a 917bp inversion of the genomic DNA sequence on the reciprocal end. One inversion breakpoint (201,692,537) was close to the primary RT breakpoint (201,692,544), supporting an origin from a single L1-induced DSB. The presence of 14bp homology (‘-14bp’) between the two breakpoints at the inversion junction supports the self-annealing step as shown in **Figure 4A**. By contrast, the absence of homology (‘0bp’) between the 3’-end of the first cDNA strand (arrowhead) and the genomic DNA breakpoint (201,691,621) supports an end-joining step instead of a template-switching mechanism. In cancer genomes, we identified three examples indicating inversion on the reciprocal ends and one example indicating inversion on the primary RT end (**Figure 4C**). Together, these observations suggest that both the primary RT end and the reciprocal end generated by ORF2p can undergo DSB end resection and remodeling to create genomic DNA inversions.

We observed inverted duplication outcomes in an experimental L1 clone and in a cancer genome. In the example from a Dox L1 clone (**Figure 4D**), the 2bp microhomology at the inversion junction supports self-annealing with a small hairpin. The example from the cancer genome (**Figure 4E**) had similar features. The small hairpins implicated in the inverted duplications contrast with the large hairpins implicated in the inversions; the different rearrangement outcomes could reflect different outcomes of small or large hairpins at DSB ends ^44,45^.

In summary, we observed L1 insertions accompanied by short deletions, inversions, or inverted duplications near the insertion site and propose that these arise from remodeling of the ORF2p-generated DSB ends prior to ligation between the primary RT and the reciprocal ends.

### Foldback rearrangements from replication/fusion of L1-induced DSB ends

The third class of rearrangements from L1-induced DSB are foldbacks. In a recent study, we demonstrated that a foldback junction can originate from a single DSB end that undergoes replication/fusion^31^ (**Figure 5A**). From 10 cancer genomes, we identified four foldback junctions containing polyadenylated L1 insertions, including two where the reciprocal breakpoint of the primary RT breakpoint was also identified at a translocation junction (**Figure 5B** and **Figure S4A**). These examples and the example shown in **Figure 1B** provide strong support for the breakage-replication/fusion mechanism. See suppl. Fig. 17 from Ref. ^31^ for examples of foldback junctions containing polyadenylated TPRT insertions identified in the experimental L1 clones^31^.

**Figure 5.**
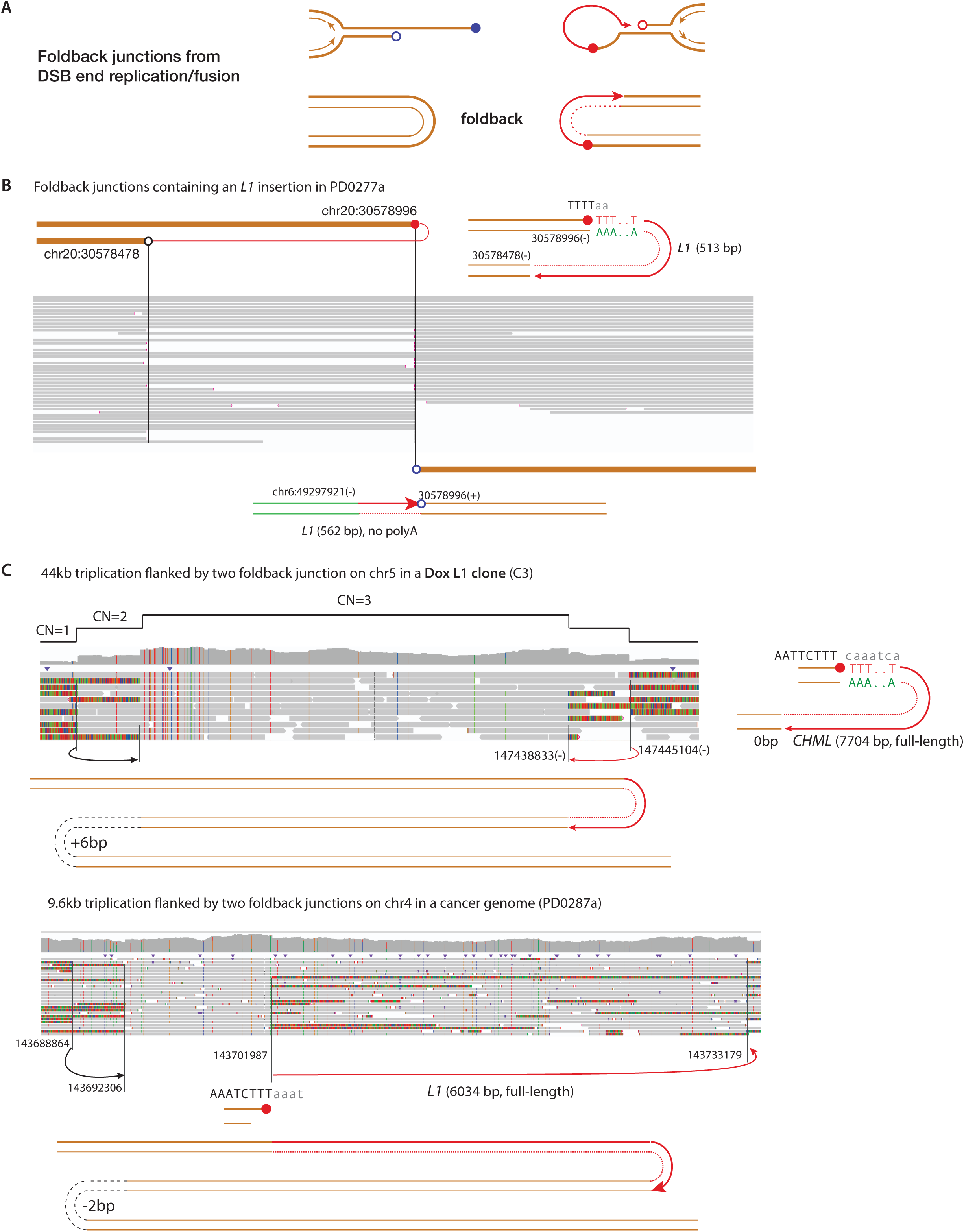
Foldback rearrangements from replication/fusion of ORF2p-induced DSB ends. **A.** Two possible foldback junctions that can form on the reciprocal end (left) or the primary RT end (right). Foldback of the primary RT end contains a TPRT insertion. **B.** An example of foldback junction with a reciprocal breakpoint in a translocation junction identified in a cancer genome. The polyadenylated insertion identifies the ancestral 3’ primary RT end (red filled circle) at chr20:30,578,996(-). The reciprocal breakpoint (blue open circle) at 30,578,996(+) is likely the ancestral 5’ reciprocal end. The primary RT end folds back to form the foldback junction after replication/fusion. The reciprocal end forms a translocation junction to chr6 that contains a non-polyadenylated L1 insertion. The breakpoint configuration is nearly identical to the example shown in **Figure 1B**. Another similar example is shown in **Figure S4A**. **C.** Two examples of DNA triplication flanked by foldbacks on both sides, one of which containing a TPRT insertion. In the top example from an experimental L1 clone, the TPRT insertion is a full-length cDNA of the *CHML* gene; in the bottom example from a cancer genome, the TPRT insertion is a full-length L1. In both **B** and **C**, the foldback junction cannot have derived from the ends of broken bridge chromosomes, as the distal sequence telomeric to the breakpoint is preserved in the same clone. The examples of foldbacks with a reciprocal breakpoint (shown in **B**, **Figure 1B**, and **Figure S4A**) can be straightforwardly explained by the model shown in **A**. The triplication outcome shown in **C** may be explained by a speculative mechanism shown in **Supplementary Figure 2**.

Foldback junctions can also originate from the reciprocal end and do not contain TPRT insertions. The inverted duplication outcome shown in **Figure 4D** and **4E** can be regarded as such instances. If the primary RT end that contains the TPRT insertion is deleted, then the only sequence feature of a foldback junction on the reciprocal end is the presence of potential ORF2p EN target sequences. In the experimental L1 clones, we identified multiple instances of foldback junctions without TPRT insertions but with breakpoints adjacent to sequences amenable to ORF2p EN cutting (**Figure S4B,C**). By contrast, no foldback rearrangement was detected in control clones (Table S3). Therefore, we expect at least a subset of these foldback junctions to have derived from the reciprocal ends generated by ORF2p.

Finally, we identified two instances of triplication flanked by two foldback junctions, one of which containing a TPRT insertion (**Figure 5C**). These outcomes recapitulate the “dup-trp-dup” copy-number pattern previously described in both congenital disorders^51^ and cancer genomes (also known as “local 2-jumps”^52^). Interestingly, in both examples observed here, the insertion sequences indicate reverse transcription of full-length mRNAs; we speculate that these triplication outcomes result from a re-replication process during which an L1-mediated foldback junction is formed by a direct ligation between an extended RT end and another genomic DNA end (**Supplementary Figure 2**).

In summary, we suggest that foldback junctions containing TPRT insertions and/or adjacent to ORF2p EN target sequences are both explained by replication/fusion of ORF2p-generated DSB ends^31^.

### Rearrangements originating from ssDNA ends due to incomplete second-strand synthesis

The rearrangement outcomes discussed above can all be explained by either remodeling (inversions or foldbacks) of L1-induced DSB ends or their ligation to distal DSB ends (translocations). For two DSB ends generated at a single TPRT target site, we expect them to be either ligated together to create an insertion junction or ligated to distal DNA ends to form two translocation junctions. Surprisingly, we identified two instances in cancer genomes where both translocation and insertion outcomes were observed at the same locus (**Figure 6**). Based on similar outcomes of insertion rearrangements (suppl. fig. 12 of Ref. ^31^), we propose that these two outcomes derive from opposite strands of the ancestral DNA with a ssDNA gap (**Figure 6A**). On the top strand, the 3’ primary RT end is extended by TPRT and ligated to the 5’-reciprocal end; this strand is converted into a dsDNA with an insertion. On the bottom strand, a ssDNA gap is formed due to incomplete second-strand synthesis and/or resection of the 5’-primary RT end; the 3’ and 5’ ends of this ssDNA gap are converted into two DSB ends by replication (i.e., ssDNA breakage-replication/fusion^31^), resulting in a deletion at or near the 3’-end of the TPRT insertion (primary RT end). If these newly generated DNA ends are ligated to each other, the outcome is a 3’-truncated insertion (without polyadenylation) with genomic DNA deletion. If the newly generated DNA ends are ligated to distal DNA ends, there will be two translocation or long-range rearrangement junctions, one of which contains a 3’-truncated TPRT insertion.

**Figure 6.**
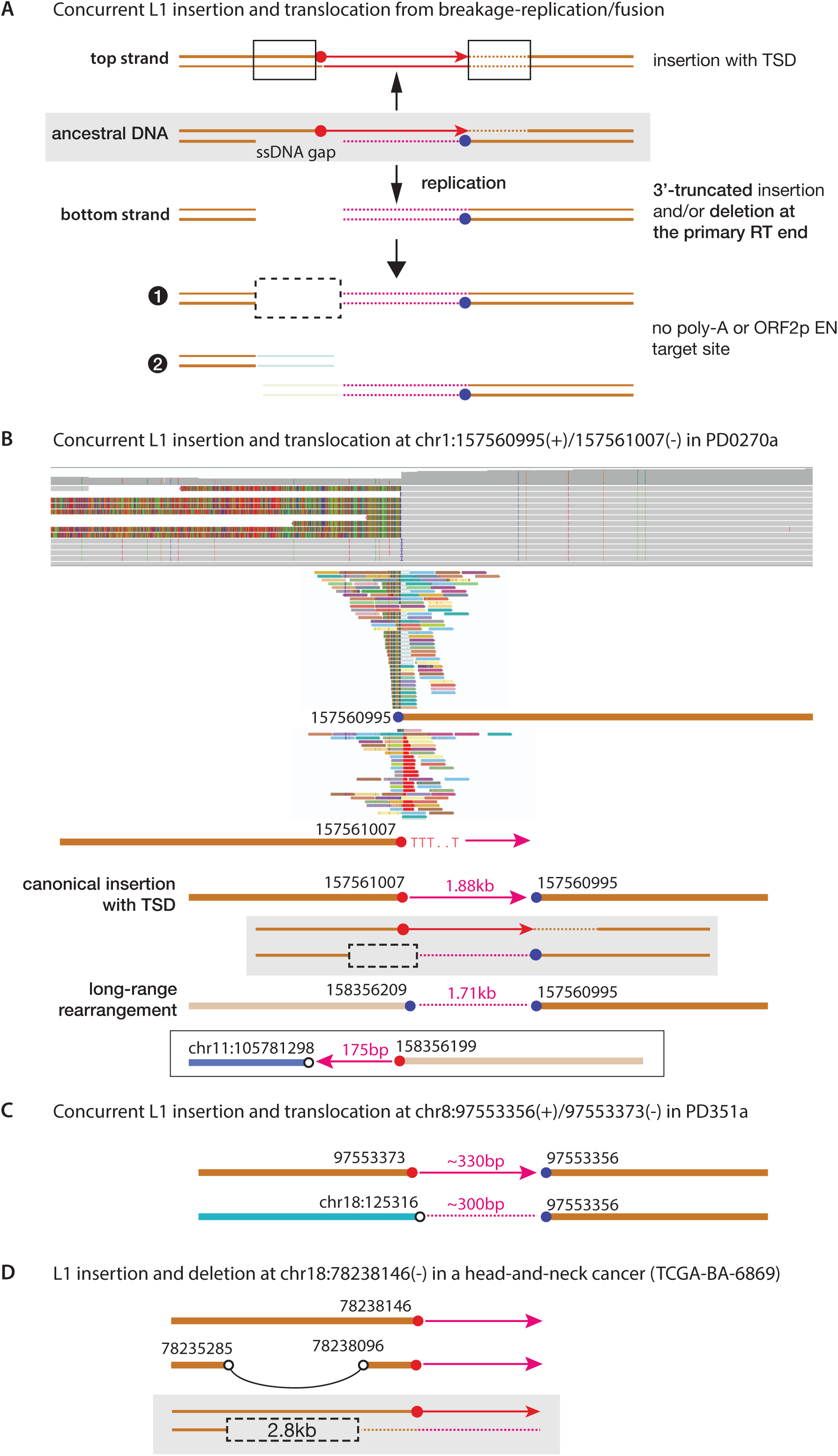
Concurrent insertion and translocation outcomes at a single ORF2p target site from incomplete second-strand synthesis. **A.** A proposed model where an incomplete TPRT event leads to two different alterations. The completely ligated top strand (first cDNA strand) of the ancestral DNA generates a TPRT insertion with all the canonical TPRT features (polyadenylation, ORF2p target sequence, TSD). A ssDNA gap in the bottom strand is converted to two dsDNA ends by replication. The ssDNA gap can arise from incomplete second-strand synthesis or 5’-resection of the primary RT end. The two dsDNA ends can be ligated together to create a 3’-truncated insertion, or ligated to other DSB ends to create a pair of translocations. In both scenarios, the canonical TPRT features (polyadenylation, ORF2p target sequence, TSD) may be lost due to the deletion. **B.** An example of concurrent L1 insertion and translocation identified in a cancer genome. The top panel shows a snapshot of long reads with either soft clipping indicating a rearrangement outcome and an insertion; the middle panels show short reads with the same soft clipped sequences on both sides. Two breakpoints indicated by each group of soft-clipped short reads (chr1:157,560,995 and 157,561,007) display a 13bp duplication. The bottom panel shows the segmental structure of DNA sequences corresponding to the insertion and rearrangement outcomes. The inferred structure of the ancestral DNA strands is shown in the middle. Note that the partner breakpoint at chr1:158,356,209 is itself derived from a DSB end that is reciprocal to a primary RT end (chr1:158,356,119) identified in a translocation junction (shown in the box). The opposite orientations of the two insertions are incompatible with homologous recombination since the two L1s are anti-parallel. **C.** Another example similar to **B** identified in the same genome. **D.** A single TPRT insertion feature preserved in two different DNA sequences in a cancer genome. Although we are unable to determine the complete junction sequence, we can identify two different genomic DNA sequences both in *cis* with the insertion, one of which containing a deletion. This observation is easily explained by a gap within the genomic DNA created by 5’-resection of the ancestral primary RT end.

In the first example shown in **Figure 6B**, we identified a 5’-truncated (1.88 kb) L1 insertion and a translocation junction with an insertion (1.71kb) with further truncation on the 3’-end at chr1:157,560,995. The presence of a translocation junction and an insertion junction at the same locus is explained by the model shown in **Figure 6A**. We further determined the translocation partner (chr1:158,356,209) derived from a reciprocal DNA end at a different TPRT site. Note the TSD feature between the reciprocal breakpoints and the primary RT breakpoints at both target sites. A similar example was found in another cancer genome (**Figure 6C**). In a third example from a different cancer genome (**Figure 6D**), we identified two different genomic DNA sequences with the same *de novo* L1 insertion, one of which showed a 2.8kb deletion near the 3’-end of the integrated L1. This observation can be explained by the same mechanism as shown in **Figure 6A** when the ssDNA gap arises from resection of the 5’-end of the primary RT end.

Together, the observations of two different rearrangement or insertion outcomes at a single site of L1 integration suggest that DNA rearrangement can also arise from ssDNA gaps due to incomplete second-strand synthesis and/or 5’-resection of the primary RT end.

### Segmental copy-number alterations from L1-induced DSB and translocations

In above, we discussed two mechanisms that can explain two classes of genomic DNA deletions at TPRT target sites (**Figure 7A**, left): (1) deletion at the 5’-end of the insertion due to processing of the reciprocal end (**Figure 4A**); and (2) deletion at the 3’-end of the insertion due to 5’-resection of the primary RT end (**Figure 6A**). For both mechanisms, the lengths of deletions should be within the size of DSB end resection (<10kb); this is consistent with deletions observed in both cancer genomes and the experimental clones (**Figure S5A**, left). We also observed small (1-10kb) tandem duplications (**Figure 7A**, middle) at a lower frequency (**Figure S5A**, right). Small tandem duplications can be regarded as large TSDs^47^. We suggest that such duplications arise from break-induced replication from the reciprocal end that is similar to the replication bypass mechanism underlying 10kb tandem duplications in BRCA deficient cells^53^. Finally, we observed two large duplications (chr13:57.27-57.35Mb in PD0270; chr1:156.1-157.2Mb in PD0277a); these duplications can reflect either a large genomic DNA insertion at an L1 insertion site (**Figure 7A**, right) or two translocations with a large overlapping duplication, which cannot be definitively resolved by shotgun sequencing. All these outcomes have only small copy-number alterations at the site of TPRT insertion and we suggest they arise from a single L1-induced DSB.

**Figure 7.**
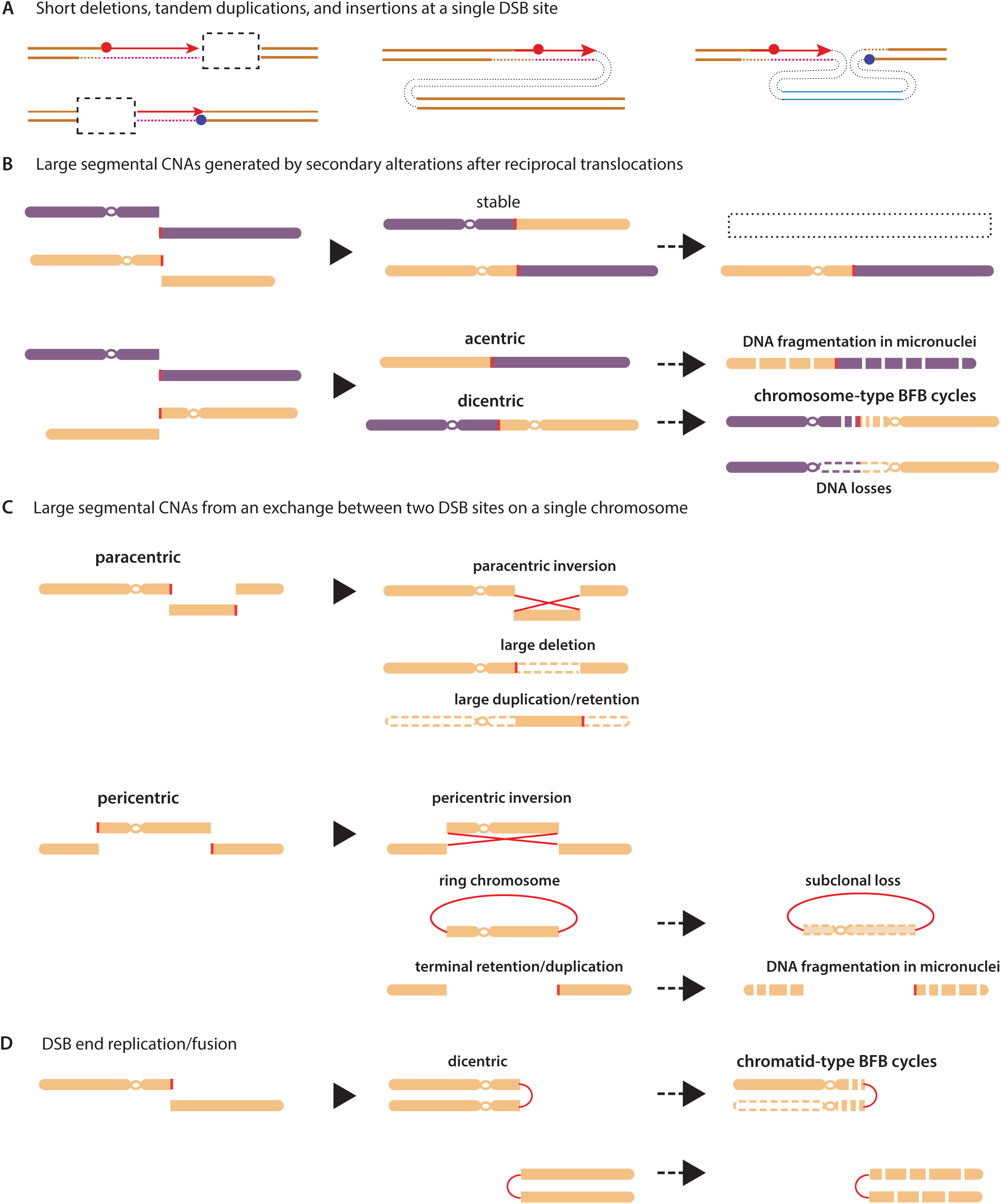
Copy-number alterations from L1-induced DNA rearrangements and chromosomal instability. **A.** Copy-number alterations and rearrangements at a single DSB site generated by L1 ORF2p. Left: small deletions near the reciprocal (cf. **Figure 4A**) or the primary RT ends (cf. **Figure 6A**) generated by ORF2p. Middle: small duplications that may result from break-induced replication from the reciprocal end. According to this model, one DNA strand remains intact and is used as template for homology-mediated DNA synthesis. Right: ectopic insertions of distal DNA segments at the ORF2p-generated DSB site. The inserted genomic DNA sequences may reach several megabases. **B.** Large segmental copy-number alterations arising downstream of reciprocal translocations generated from DSB sites on two chromosomes. Top: One of two stable chromosomes generated by the translocations is deleted (or duplicated, not shown), creating terminal deletion (orange chromatid) and retention (purple chromatid). As the remaining derivative chromosome is stable, there is no segmental copy-number variation in the retained segments. Examples indicating such a process are shown in **Figure S5B**, **S5C** (from experimental L1 clones) and **S5D** (from a cancer genome). Bottom: If the reciprocal translocations generate an acentric and a dicentric chromosome, both unstable chromosomes will acquire further copy-number alterations and rearrangements. Examples indicating this scenario are shown in **Figure S6** (all from experimental L1 clones). **C.** Large segmental copy-number alterations arising downstream of intra-chromosomal rearrangements generated from two DSB sites. Top: Two possible outcomes of large segmental CNA generation from an exchange between two pairs of DSB ends on a single chromosome arm. One outcome is a large inversion with no CNA, as shown in **Figure S3A**. Another outcome is a large deletion with examples shown in **Figure S5E** (from an experimental L1 clone) and **S5F** (from a cancer genome). The excised acentric fragment may be integrated as an ectopic insertion (e.g., as shown in panel **A**). Bottom: When two DSB sites occur on opposite arms, an additional outcome is a ring chromosome. One such example is shown in **Figure S5G** (from an experimental L1 clone). **D.** Unstable chromosomes can also arise from replication/fusion of either DSB end generated by ORF2p at a single site.

We suggest that large (1Mb or above) CNAs with L1-mediated rearrangements arise from two DSBs (**Figure 7B,C**). The simplest mechanism that can lead to large terminal deletions or duplications is by deletion or duplication of one of two translocation chromosomes (**Figure 7B**, top). Selected examples from both experimental L1 clones and cancer genomes are shown in **Figure S5B-D**. When two pairs of DSB ends are generated on a single chromosome, a reciprocal exchange between these ends can generate a large deletion (**Figure S5E,F**) or a ring chromosome with two terminal deletions (**Figure S5G**). The excised acentric fragments may be translocated or inserted into other loci, creating duplications of a large internal or terminal segments. All these copy-number outcomes arise directly from stable derivative chromosomes generated by L1-mediated translocations.

Large CNAs can also arise from the downstream evolution of unstable chromosomes generated by L1-mediated translocations. As mentioned above, a simple reciprocal exchange between DSB ends on two chromosomes can generate two unstable chromosomes, one dicentric and the other acentric (**Figure 7B**, bottom). Dicentric chromosomes can form chromosome bridges and acentric chromosomes can form micronuclei, both of which can cause chromosome fragmentation and generate chromothripsis^34,54–56^. Both outcomes were recapitulated in our experimental L1 clones and single cells (**Figure S6**). In comparison to CNAs generated by gain or loss of stable chromosomes, the evolution of dicentric or acentric chromosomes will often generate complex rearrangements (**Figure S6A**) or copy-number variation on the retained segments (e.g., **Figure S6B,C**). By contrast, stable translocated chromosomes display constant DNA copy number across the retained segments.

With segmental copy-number losses or deletions, the breakpoints derived from ancestral L1-induced DSB ends may be lost during clonal expansion. In the example shown in **Figure S7A** from a cancer genome, we inferred that the 10q segment (59Mb-qter) was retained in a stable derivative chromosome with an L1-mediated translocation junction (red vertical line); the complex amplifications within the remaining region of chr10 (0-55Mb) may have originated from an unstable chromosome generated by translocations involving the reciprocal end generated by TPRT.

Downstream evolution of unstable chromosomes generated by L1-mediated translocation can also result in deletion of TPRT insertion junctions. Two examples of dicentric chromosomes that may have resulted from fusions between the reciprocal ends generated during TPRT are shown in **Supplementary Fig. 3**. Notably, in the experimental samples, we observed large CNAs only in the L1 clones (**Supplementary Fig. 4**) but not in control clones. In L1 but not control clones, we also observed many instances of terminal deletions that were accompanied by variable segmental losses between the deletion breakpoint and the centromere (“sloping copy-number variation”) (**Supplementary Fig. 5**); these patterns are consistent with the breakage-fusion-bridge cycles of dicentric chromosomes^34,57,58^. It is plausible that these alterations were instigated by L1-induced rearrangements but the TPRT insertion features were lost during downstream evolution.

Finally, we observed multiple instances of chromothripsis in the experimental L1 clones (**Figure S7B, S7C**; **Supplementary Fig. 6**) where TPRT insertions (both polyadenylated and non-polyadenylated/3’-truncated) were identified in a subset of rearrangement junctions. Rearrangement junctions with 3’-truncated L1 insertions were also observed in cancer genomes. Although these junctions were previously thought to indicate ORF2p EN-independent reverse transcription^5^, we suggest that at least some of these junctions arise from incomplete 2^nd^ strand synthesis during TPRT as shown in **Figure 6A**.

In summary, L1-induced chromosome rearrangements and translocations can both directly generate large CNAs and fuel subsequent copy-number evolution by generating unstable chromosomes that can acquire complex rearrangements including chromothripsis.

## DISCUSSION

Since the discovery of DNA transposition by McClintock^58,59^, transposition has been recognized as a major driver of genome evolution^60,61^. While the most common outcomes of transposition are transposon insertions, transposon activity can cause DNA damage^28^ and has been associated with profound karyotypic changes in speciation^62,63^ and in human cancer genomes^27,30^. However, the full spectrum of genomic changes caused by L1 retrotransposition remains unknown. Here, we provide, to our knowledge, the first comprehensive characterization of L1-induced rearrangements in both experimental models and primary cancers by a combination of short- and long-read sequencing. Observations from our experimental systems after a short period of potent L1 expression recapitulate all classes of retrotransposition-associated rearrangements in cancer genomes. These alterations can be explained by illegitimate recombination, remodeling, and replication of DNA ends generated by L1 ORF2p, or the downstream evolution of unstable chromosomes generated by L1-induced rearrangements. Our findings significantly expand the repertoire of L1-induced genomic alterations beyond L1-mediated deletions^3,46–48^, and provide new insights into the molecular processes of retrotransposition.

### Translocations from illegitimate recombination between L1-induced DSB ends

The first class of L1-induced rearrangements are translocations or rearrangements arising from a reciprocal exchange between L1-induced dsDNA ends. Such events can lead to intra-chromosomal inversions, reciprocal translocations between two chromosomes, or chromoplexy (**Figure 3** and **S3**). The similarity between these outcomes and those generated by site-specific DSBs^64^ or after genome-wide DNA damage by X-ray irradiation^65,66^ suggests non-homologous end-joining between reciprocal DSB ends as the single underlying mechanism. Notably, based on the reduction in clonogenicity (∼50%) and the number of copy-number transitions in the progeny clones (1-2 per genome), we estimate that the degree of DNA damage generated by L1 induction in our experimental system is comparable to 2Gy X-ray radiation^66^.

The inference that the ancestral DSB ends in such translocations are generated by L1 ORF2p is established by three sequence features of the translocation junctions: (1) polyadenylated insertions that identify the ancestral 3’-end of TPRT; (2) ORF2p EN target sequences at the site of TPRT; and (3) short duplications between the primary RT and the reciprocal breakpoints resembling the TSD feature of canonical insertions, indicating two ancestral DNA ends with short 3’-overhangs (‘sticky’ ends).

The alternative explanation that such translocations arise from non-allelic homologous recombination (NAHR) between two pre-existing L1’s (either endogenous or *de novo*) can be excluded based on three observations. First, we do not observe extensive homology at the translocation junctions as expected for NAHR. Second, NAHR between two L1 sequences should only occur between two L1’s with the same orientation; by contrast, many translocation junctions are between two L1’s with opposite orientations. Moreover, in our Tet-On L1 expression system, many translocation junctions contain one L1 insertion and one cDNA insertion with no sequence homology. Third, for NAHR between two L1 sequences, we expect to see partial L1 sequences at both genomic DNA breakpoints from each site; this is not observed for L1 insertions. In one exceptional case, we observe two truncated cDNA insertions at both breakpoints (**Figure S3C**); however, the inverted orientations of the two cDNA fragments suggest twin priming^43^. Finally, we also observe reciprocal translocations between reciprocal breakpoints from an L1 integration site and breakpoints without TPRT signatures, indicating recombination between L1-induced DSB ends and L1-independent DSB ends^42^. Together, these observations suggest that L1-mediated translocations arise from recombination between L1-induced DSB ends, but not homologous recombination between pre-existing L1 sequences.

### Short inversions, inverted duplications, and foldback rearrangements from remodeling of L1-induced DSB ends

Unligated DSB ends can undergo 5’ resection^44,67^ to have 3’-overhangs. Self-annealing of the 3’-overhang can give rise to inversions or inverted duplications^45,68^. We suggest that this mechanism underlies L1 insertions with local genomic DNA inversion and inverted duplication that are observed in both cancer genomes and our experimental systems (**Figure 4**). Unligated DSB ends that persist into S/G2 can also form foldback junctions by replication/fusion^31,69^. The breakage-replication/fusion mechanism (**Figure 5A**) provides a simple explanation of foldback junctions containing TPRT insertions (**Figure 1B, 5B, S4A**).

### Mechanistic implications of L1-induced rearrangements on DNA breakage and ligation during retrotransposition

The translocation, inversion, and foldback outcomes of L1-induced rearrangements all indicate untethered DSB ends generated during TPRT. These observations further provide insights into the mechanisms of DNA breakage and ligation during TPRT *in vivo* that complement biochemical analyses of ORF2p.

First, the observations of short duplications between reciprocal translocation breakpoints (**Figure 3**) resembling the TSD feature of canonical insertions indicate a single ancestral configuration of staggered breaks on opposite strands. The TSD feature is also preserved in the inversion (**Figure 4C**) and inverted duplication (**Figure 4D,E**) outcomes. These observations suggest that the staggered break end configuration reflect a specific positional constraint of the second-strand nick relative to the first-strand nick (primary RT end). This inference is consistent with our recent finding that ORF2p EN robustly cleaves second-strand DNA based on structural distortion of the helix^9^ but not sequence specificity. Notably, reciprocal translocations generated by other processes of DSB, including designer nucleases^64^ or radiation^66^ rarely display the TSD feature, further indicating the staggered configuration as a specific feature of ORF2p-generated DSB ends. Whether this feature is solely due to ORF2p or also involves host proteins will require additional studies.

Second, the observations of inversions or inverted duplications suggest significant 5’-resection of ORF2p-generated DSB ends. The 3’-ssDNA overhang may undergo deletion due to flap removal during microhomology-mediated end-joining^64,70^ that abolishes the TSD feature, even though the ancestral DSB ends generated by ORF2p have the staggered end configuration.

Third, the observations of concurrent insertion and translocation outcomes at a single target site (**Figure 6B,C**) and genomic DNA deletions at the 3’-end of TPRT insertions (**Figure 6D**) suggest the presence of large ssDNA gaps due to incomplete second-strand synthesis or 5’ resection of the primary RT end (**Figure 6A**). Replication/fusion of the ssDNA gap generates 3’-truncated (i.e., non-polyadenylated) insertions or insertion-mediated rearrangements. Previously, 3’-truncated insertions have been ascribed to EN-independent functions of ORF2p RT acting on a preexisting 3’-OH DNA end^5^. Our data definitively show such outcomes can result from an ORF2p EN-mediated TPRT event.

Finally, the observations of triplications flanked by TPRT-mediated foldbacks (**Figure 5C**) suggest TPRT after only single-strand nicking. This is because if ORF2p nicks both DNA strands, there cannot be an intact copy of the DNA sequence across the breakage site that is retained in the triplication. Therefore, the activity of ORF2p EN is not limited to target sites with discontinuities on the opposite strand^10,13^. Moreover, as the generation of foldbacks involves a replisome passing through an unligated DSB end, the observations of TPRT-mediated foldbacks therefore indicate that ORF2p can generate double-stranded primary RT ends prior to DNA replication.

### DNA copy-number alterations from L1-induced translocations and rearrangements

L1-induced translocations and rearrangements can lead to several classes of copy-number alterations of different sizes (**Figure 7**). The most common alterations are L1-mediated small deletions; the sizes of such deletions are usually less than 10kb and within the size range of 3’ flaps of resected DSB ends. Less common are L1-mediated small duplications (1-10kb), which can arise from break-induced replication^53^ initiated from the reciprocal end. We suggest that large segmental CNAs (>1Mb) arise from long-range intra-chromosomal rearrangements or inter-chromosomal translocations followed by deletion or duplication of one or both translocated chromosomes. Finally, L1-induced translocations can create acentric, dicentric, and ring chromosomes that are mitotically unstable. These unstable chromosomes can undergo chromosome fragmentation or breakage-fusion-bridge (BFB) cycles to create complex rearrangements and copy-number variation. It is noteworthy that McClintock’s original discovery of DNA transposition was based on the appearance or disappearance of chromosomal instability, including large terminal deletions and BFB cycles, that marks DNA breakage due to self-excision of DNA transposons.

### The impact of L1 retrotransposition on somatic genome evolution and tumorigenesis

Our findings suggest that L1 expression in cancer^18^ or pre-cancer cells can directly contribute to the acquisition of large-scale chromosomal rearrangements and copy-number alterations. L1-induced genome instability is initiated by L1-induced DSB and compounded by the evolution of unstable chromosomes generated by L1-induced translocations or rearrangements during clonal expansion. As rearrangements arising during the downstream evolution are not directly caused by TPRT, their breakpoints do not have features of TPRT; therefore, the overall impact of L1 retrotransposition on genome evolution exceeds insertion-mediated rearrangements. This conclusion is supported by the observation of long-range rearrangements without TPRT features in experimental L1 clones in comparison to no such alterations in the control clones. It can also explain the positive correlation between L1 insertions and chromosome rearrangements in p53-mutated cancers^27,71^. Finally, it suggests L1 dysregulation in cancer and cancer precursors^72–74^ as a trigger of cancer genome instability and complexity.

The generation of L1-induced translocations or rearrangements requires two or more pairs of DSB ends. An outstanding question is whether and how frequently L1 ORF2p can generate two or more DSBs in a single nucleus. In our experimental systems with transient but potent L1 expression, we observe a similar number of insertion and translocation outcomes. By contrast, the cancer genomes contain a much higher burden of L1 insertions than L1-mediated translocations. We speculate that L1-mediated translocations may result from episodes of high L1 activity that creates many DSB ends all-at-once, whereas low or modest L1 activity primarily gives rise to L1 insertions. The preponderance of L1 insertions may then reflect tolerance of chronic L1 activation but not sustained high L1 activity. Notably, in cancer genomes, L1-mediated translocations occur both before and after large segmental or chromosomal duplications, suggesting that the triggers of L1-mediated translocations are active throughout the course of cancer genome evolution. Given that potent L1 expression significantly reduces cell fitness and clonogenicity^29^, harnessing mechanisms that cause L1 hyperactivity may provide new strategies for tumor-specific genotoxic therapy.

In summary, our study suggests L1 retrotransposition as a potent instigator of genome instability that can lead to complex rearrangements and copy-number variation, underscoring an unappreciated connection between L1 activity and genome complexity in cancer cells.

## Data Availability

All sequencing data of RPE-1 cells and RPE-1 clones generated in this study have been deposited to the Sequencing Read Archive (SRA) under BioProject PRJNA1197453.

## AUTHOR CONTRIBUTIONS

C.-Z.Z and K.H.B. conceived the project. C.M-D, K.H.B., and C-Z.Z. designed the experiments. C.M.D. performed all the experiments and experimental data analyses with help from P.S. and J.C.K. C.M-D. and J.A.K. performed the DNA copy-number analysis of low-pass whole-genome sequencing data. X.Z. and J.A.K. performed the differential expression analysis and the analysis of insertions. S.T. processed PacBio Hi-Fi reads of the experimental L1 clones and both long-and short-read sequencing of the cancer genomes. M.L. performed the haplotype-specific DNA copy-number analysis of cancer genomes. S. Z. and J.M.C.T. performed the sequencing of cancer genomes. J.W. assisted in single-cell whole-genome sequencing data generation. C.-Z.Z performed the analysis of L1-mediated genomic rearrangements and copy-number alterations in both experimental and cancer genomes with help from X.Z., J.A.K., and S.T. K.H.B. supervised the experimental analyses. C.-Z.Z. supervised the genomic analyses. C.M-D. K.H.B and C-Z.Z. wrote the manuscript with input from all authors.

## FUNDING SOURCES

K.H.B., E.A.L and C-Z. Z. are supported by the National Cancer Institute (R01CA276112) K.H.B. is supported by the National Institutes of Health (R01CA240816, R01CA289390, UG3NS132127). E.A.L was supported by the National Institute of Health (NIH) (DP2 AG072437) and the Suh Kyungbae Foundation. K.H.B. and C-Z. Z. are supported by the Dana-Farber Cancer Institute (Innovations Research Fund). J.A.K. was supported by a postdoctoral fellowship from the American Cancer Society (PF-22–123–01-DMC). C.M-D is a Fellow of The Jane Coffin Childs Fund for Medical Research.

## METHODS

### Cancer genome analyses

We obtained the following whole-genome sequencing (WGS) data for the analysis of retrotransposition-related rearrangements and copy-number alterations in cancer genomes. 1. Short-read WGS data of 13 cancers from The Cancer Genome Atlas that have the highest burdens of somatic L1 insertions, and their matching germline reference samples, all downloaded from the National Cancer Institute Genomic Data Commons. 2. Short- and long-read whole-genome sequencing data of 10 cancer genomes published in a recent study. We performed haplotype-specific copy-number analysis and identified somatic retrotransposition events from published data that coincide with copy-number transitions.

### Haplotype-specific copy-number calculation

Haplotype-specific DNA copy number was calculated following the same strategy as described previously. Briefly, we first calculated haplotype-phased allelic coverage in 10kb bins from the total sequence coverage and the phased allelic fraction; we then performed long-range haplotype phasing in regions of allelic balance. The normalized, haplotype-specific DNA copy number was then used to identify sites of copy-number transitions co-localized with somatic L1 integrations reported in the previous studies.

### Experimental systems of L1 expression

#### Cell culture

Cells were cultured at 37°C in 5% CO2 atmosphere with 100% humidity. hTERT-immortalized, p53-/- RPE-1 cells and U2OS Flp-In T-REx cells were grown in DMEM/F12 (1:1) or DMEM respectively, and supplemented with 10% FBS, 100 IU/ml penicillin, and 100 μg/ml streptomycin. For cell lines containing doxycycline-inducible transgene constructs, tetracycline-free FBS (Takara) was used in all culture media, and 1 μg/ml of doxycycline was used to induce transgene expression. U2OS cells were provided by Dr. Jeremy Stark (1), and RPE-1 cells were provided by Dr. David Pellman (2).

#### Tet-On L1 expression system

We used several plasmid constructs to induce the expression of a codon-optimized sequence of human L1 (ORFeus) in cells. For the Tet-On expression system in RPE-1 cells, we exploited the Sleeping Beauty DNA transposon system to stably deliver a Tet-On L1 (ORFeus) cassette, which we previously validated (3). For this, we transfected hTERT RPE-1 p53-/- cells using Viafect (Progema) with an expression vector for the Sleepy Beauty transposase (pCMV(CAT)T7-SB100), and the donor plasmid containing Sleepy Beauty inverted terminal repeats flanking the inducible Tet-On L1 cassette, and a constitutively expressing rtTA-T2A-NeoR cassette (pCMD20D). Cells were selected with G418, and single cells were sorted to generate a monoclonal cell line (Tet-On L1 RPE-1 p53-/- cells), which we confirmed via immunoblotting of the L1-encoded proteins. Similarly, we also generated a monoclonal cell line containing a Tet-On Luciferase cassette (pCMD26A, Tet-On Luc RPE-1 p53-/-). For transient expression of L1 in RPE-1 cells or U2OS cells, we used a pCEP4 episomal expression vector that was modified to contain a Puromycin resistance gene to express under a CMV promoter a codon-optimized sequence of human L1 containing a GFP reporter for retrotransposition (pMT527, pCEP4 L1-ORFeus), which we previously validated (4). We also cloned retrotransposition mutant versions of the L1 reporter plasmid containing mutations in the encoded ORF2p, including a reverse transcriptase mutant (pLD631, D702Y) (4) and two endonuclease mutants (pCMD104A, E43A:D145A & pCMD103A, D205G:H230A). To establish the Tet-On expression system in U2OS Flp-In T-REx cells, we removed the GFP cassette from pcDNA5 FRT/TO GFP (Addgene #19444) and cloned in our L1 expression cassette derived from our pCEP4 vectors aforementioned: pCMD111az (pcDNA5 FRT/TO L1-ORFeus), pCMD112az (pcDNA5 FRT/TO RTmutant D702Y), pCMD113az (pcDNA5 FRT/TO L1-ORFeus ENmutant D205:D145A) and pCMD114az (pcDNA5 FRT/TO L1-ORFeus ENmutant E43A:D145A). These Tet-On L1 expression plasmids were integrated into U20S Flp-In T-Rex cells by co-transfection with the PGK-Flp recombinase vector using FugeneHD (Promega), as previously described (1). Integrated clones were selected using hygromycin (0.2 ug/uL) and subsequently screened by immunoblotting for both L1-encoded proteins.

#### L1-GFP reporter system

We performed the L1 GFP reporter assay for retrotransposition in RPE-1 p53-/- cells as previously described (3, 5). Briefly, we seeded 0.8 x10^5^ cells in 6-well plates (day 1, d 1). The following day we transfected each well using Viafect (Promega) with 1 ug of pCEP4-Puromycin L1 GFP reporter plasmid (d 2): pMT527 (WT), pLD631 (RTmutant-D702Y), pCMD103A (ENmutant-D205G:H230A) or pCMD104A (ENmutant-E43A:D145A). A pCEP4-Puromycin vector expressing a GFP cassette (MT498, pCEP4-GFP) was used to monitor transfection efficiency by assaying the percentage of GFP+ cells on d 4 by flow cytometry (BD LSRFortessa). Media was changed 12 hours after transfection (d 3). Two days after transfection (d 4), the media for cells transfected with the L1 reporter plasmids were supplemented with 5 ug/mL of Puromycin. Cells were then incubated until d 7 when cells were collected and assayed for the percentage of GFP+ cells by flow cytometry (BD LSRFortessa). Singlets were gated on side-scatter versus forward scatter and GFP+ cells were gated on GFP versus autofluorescence (PE). Cells with autofluorescence were detected on a diagonal line, whereas cells showing increased green fluorescence (GFP+) were gated above the autofluorescence diagonal line. We normalized the percentage of GFP+ cells from the cells transfected with the L1 reporter assay to the percentage of GFP+ cells from cells transfected with the GFP plasmid.

### Molecular and cell biological analyses of experimentally induced L1 expression

#### Induction of L1 expression in the Tet-On system

Tet-On L1 p53-/- RPE-1 cells were exposed to Dox for five days to induce L1 expression. Tet-On Luc (also p53-/-) RPE-1 with the same treatment and Tet-On L1 p53-/- RPE-1 cells treated with DMSO for five days were used as control.

#### Immunoblotting

For Tet-On L1 or Luc RPE-1 p53-/- cells, cells were treated with DMSO or Dox (1 µg/mL) for 5 d and collected for protein extraction. As a control, parental RPE-1 p53-/- cells were treated with DMSO, or 1 µM mitomycin C (MMC, SC-3514) for 48 h. For U2OS Flp-In T-REx cells transfected with L1-expressing pCEP4 vectors, 2x10^5 cells were transfected with 1 ug of plasmid and collected two days after for protein extraction. For Tet-On L1 U20S cells, cells were treated with Dox (1 µg/mL) and collected three days later. Protein was extracted from cells using radioimmunoprecipitation assay buffer (Boston BioProducts BP-115) supplemented with protease and phosphatase inhibitors (Cell Signaling, 5872S). Gel electrophoresis was performed on protein extracts using 4 to 20% Mini-PROTEAN TGX gels (Bio-Rad, 456-1095). Proteins were then transferred to low fluorescence polyvinylidene difluoride membranes using Trans-Blot Turbo (Bio-Rad). Membranes were blocked using EveryBlot Blocking Buffer (Bio-Rad, 12010020) or Intercept Blocking Buffer (Li-COR, 927-60001), and probed with primary antibodies for pKAP1-S824 (Abcam, ab84077), KAP1 (Abcam, ab22553), RAD50-S635 (Cell Signaling, 14223S), RAD50 (Cell Signaling, 3427T), ORF2p (abcam, ab263071), ORF1p (Millipore Sigma, MABC1152), beta-tubulin (Cell Signaling, 2128S), gH2AX (Cell Signaling, 2577S), H3 (abcam, ab1791), CHK1 (Cell Signaling, 2360S), pCHK1-S345 (Cell Signaling, 2348S), RPA32 (Bethyl, A300-244), pRPA32-S4/S8 (Bethyl, A300-245), or pRPA32-S33 (Bethyl, A300-246), followed by secondary antibodies (IRDye 800CW goat anti-mouse IgG, 925-32210; IRDye 680RD goat anti-rabbit IgG, 925-68071; and anti-rabbit IgG horseradish peroxidase (HRP), 7074S). ECL substrate (Thermo Scientific, 34580) was used to develop HRP signals. Immunoblotting signals were detected using the ChemiDoc imaging system (Bio-Rad).

#### Immunofluorescence

For Tet-On L1 cell lines, cells were seeded on #1.5 coverslips and treated with DMSO or Dox (1 ug/uL) for five days (RPE-1 cells) or three days (U2OS cells). U2OS cells were washed with PBS and treated with pre-extraction (20 mM HEPES, 50 mM NaCl, 1 mM EDTA, 3 mM MgCl2, 300 mM sucrose, 0.25% Triton-X 100) prior to fixation. RPE-1 cells were washed with PBS and immediately fixed with 4% paraformaldehyde for 20 minutes at room temperature. Cells were then washed with 0.1 M glycine followed by PBS prior to permeabilization with 0.5% Triton X-100 in PBS for 15 minutes. Cells were washed with PBS three times and then blocked with 3% BSA in PBS for 1 hour. Cells were washed two times with PBS, followed by a 1-hour incubation with primary antibodies: γH2AX (Cell Signaling, 2577S) or 53BP1 (Cell Signaling, NB-100-904). Cells were washed three times with 0.05% Triton X-100 in PBS. Species-specific secondary antibodies were added to the cells for 1 hour followed by three washes with 0.05% Triton X-100 in PBS. 2.5 μg/mL Hoechst 33342 in PBS was added to the cells for 10 minutes. After three washes with PBS, Prolong Diamond Antifade (Life Technologies, P36961) was used for mounting the samples on glass slides. Imaging was performed on a Leica THUNDER Imager at 40X magnification using the Leica LAS X software. Quantification of γH2AX or 53BP1 nuclear foci per cell and the frequency of cells with micronuclei was performed using ImageJ.

#### Cell survival analysis

Tet-On L1 or Tet-On Luc RPE-1 p53-/- cell lines or Tet-on L1 U20S cell lines (1x10^3^ cells) were seeded in 48-well plates in media containing DMSO as control or varying concentration of Dox as indicated in triplicates. After eight days, the plates were washed with PBS and fixed with 0.5% crystal violet, 20% methanol solution for 20 mins. Plates were rinsed with H2O and air-dried and then imaged on a scanner. Once fully dried, 100 μL of methanol was added to each well to dissolve the crystal violet, and 90% of the solution was transferred to a clear 96-well plate to measure the absorbance at 570 nm from each treatment. Absorbance from the DMSO treatment per cell line was set to 1 and the rest of the absorbance measurement under Dox treatment was normalized to DMSO. This protocol was adapted from (6).

### Transcriptome analysis of experimentally induced L1 expression

Total RNA was purified from cells with either Tet-On L1 or Tet-On Luc after five days Dox treatment using RNAeasy Mini Kit including DNase treatment. Libraries were prepared using Roche Kapa mRNA HyperPrep strand-specific sample preparation kits from 200 ng of purified total RNA according to the manufacturer’s protocol on a Beckman Coulter Biomek i7. The finished dsDNA libraries were quantified by Qubit fluorometer and Agilent TapeStation 4200. Uniquely dual indexed libraries were pooled in an equimolar ratio and shallowly sequenced on an Illumina MiSeq to further evaluate library quality and pool balance. The final pool was sequenced on an Illumina NovaSeq X Plus Instrument to generate 40 million 150bp read pairs per library at the Dana-Farber Cancer Institute Molecular Biology Core Facilities.

Sequencing reads were aligned to human genome reference (GRCh38) using STAR v2.7.11b (7) with the parameter “--outSAMstrandField intronMotif”. Gene-level and transcript-level read counts were obtained using Stringtie (v2.2.3). DESeq2 (1.44.0) was used to perform differential expression analysis between Tet-On LINE-1 RPE-1 p53-/- and Tet-On Luc RPE-1 p53-/- cells. Differentially expressed genes were defined as those with an adjusted *p*-value < 0.05. Genes with fewer than 10 read counts in five or more samples (including replicates) were excluded from analysis. Gene function enrichment analysis was performed using DAVID, supplying differentially expressed genes as defined above. Volcano plots and violin plots were generated using the results from DESeq2 and DAVID.

### Generation of whole-genome sequencing data after experimental L1 expression

#### Generation of single cells and single-cell progeny clones

Cells with induced L1 expression (Tet-On L1 under 5 days of Dox treatment) and control (Tet-On L1 with 5 days of DMSO treatment) were sorted into 96-well plates. To generate whole-genome libraries from single cells, the sorted cells were immediately lysed and underwent whole-genome amplification using the REPLI-g Single Cell Kit (Qiagen, 150345), following a previously described protocol (8, 9). The amplified DNA was purified using ethanol precipitation, quantified using Qubit, and underwent library construction. 64 single cells with L1 induction and 32 control single cells (treated with DMSO) were sent for low-pass whole-genome sequencing (0.1x median depth). 78 samples (26 Dox treated; 52 DMSO treated) with uniform coverage were analyzed for large copy-number alterations. We further generated deep whole-genome sequencing data (30x median depth) on 28 cells with L1 induction and 12 control cells (including all with detectable large segmental copy-number alterations from the low-pass data).

To generate single-cell derived clones, single cells (with or without L1 induction) were sorted into 96-well plates containing media with 20% FBS. Ten plates were collected from each treatment, and the number of progeny clones was recorded to determine the clonogenicity of cells after exposure to L1 expression (**Figure 1F**). Progeny clones were expanded in 6-well plates until confluency to generate a frozen vial. Cell pellets were used for genomic DNA extraction using PureLink Genomic DNA kit (Invitrogen) and for library construction. 60 Dox clones (derived from cells with L1 induction) and 32 control clones (derived from cells treated with DMSO) underwent low-pass whole-genome sequencing. 31 Dox clones and 10 control clones were sent for deep whole-genome sequencing (20x median depth). For 25 Dox clones and one control clones, we further generated PacBio long-read sequencing data (15x median depth).

Finally, clones were also derived from GFP(+) and GFP(-) RPE-1 cells with transient expression of the L1 GFP reporter as described above (seven days of L1 GFP expression; three independent experiments). A total of 62 clones derived from GFP(+) cells and 30 clones derived from GFP(-) cells underwent low-pass whole-genome sequencing. 29 GFP(+) clones and 5 GFP(-) clones were sent for deep whole-genome sequencing (20X median depth). For 13 GFP(+) clones, we generated PacBio long-read sequencing data (15x median depth).

#### Shotgun whole-genome sequencing data generation

200 ng of amplified gDNA from single cell samples or gDNA from progeny clones was fragmented to ∼ 500 bp on a Covaris R220 instrument using a 96 microTUBE plate (Covaris, 520078). DNA libraries were prepared using reagents from LTP Library Preparation Kit (KAPA, KK8232) for multiplexed next-generation sequencing using Unique Dual-Indexed Adapters (KK8726). Finished libraries were quantified by Qubit fluorometer, and the fragment size distribution was evaluated by Agilent Bioanalyzer 2100 or Agilent TapeStation 4200. DNA libraries were pooled from each experimental condition and subjected to low-pass whole-genome sequencing (∼0.1x mean coverage) on the MiSeq (Illumina) platform with paired-end 150bp reads to assess library quality and to estimate haplotype DNA copy-number for identifying samples with genomic alterations. Selected samples with DNA copy-number alterations were subsequently selected for deep sequencing (20x mean coverage) on the NovaSeq S4 (Illumina) platform with paired-end 150bp reads. Cells with Tet-On L1: 12 control single cells, 28 single cells with L1 induction, 10 progeny clones derived from control cells, 31 progeny clones derived from cells with L1 induction. Cells with L1 GFP reporter, 5 progeny clones derived from GFP(-) cells and 29 GFP (+) progeny clones derived from GFP(+) cells.

#### PacBio HiFi long-read sequencing data generation

To construct PacBio long-read sequencing libraries, high molecular weight (HMW) genomic DNA was first purified using the MagAttract HMW DNA kit (Qiagen) or PureLink Genomic DNA kit (Invitrogen). At least 4 μg HMW genomic DNA (> 50% of fragments ≥ 40 kb) was sheared to ∼15 kb using the Megaruptor 3 (B06010003; Diagenode), followed by DNA repair and ligation of PacBio adapters using the SMRTbell Prep Kit 3.0 (102-141-700). Each library was subsequently size-selected for 10 kb ± 20% using the PippinHT with 0.75% agarose cassettes (Sage Science). After quantification with the Lunatic (Unchained Labs), libraries were diluted to 250 pM per single molecule, real-time (SMRT) cell, hybridized with PacBio standard sequencing primer, and bound with SMRT sequencing polymerase using the Revio polymerase kit (102-739-100).

Long-read sequencing was performed on the Revio instrument using 25M SMRT Cells (102-202-200) and Revio Sequencing Plate (102-587-400), with a 2-hour pre-extension time and 24-hour movie time per SMRT cell. Quality filtering, base calling, and adapter marking were done automatically on the Revio instrument. Error correction for reads generated in circular consensus sequencing (CCS) mode was performed on-board the PacBio Revio with the vendor’s ccs software (https://github.com/PacificBiosciences/pbccs) and with the following parameters:

~~~
--all --subread-fallback --num-threads 232 \
–streamed <MOVIE_NAME>.consensusreadset.xml \
--bam <MOVIE_NAME>.reads.bam
~~~

With these settings, all reads from the instrument (including those failing error correction) were presented in a single BAM file for downstream analysis. Multiplexed, barcoded libraries were demultiplexed automatically on-instrument. CCS reads for each barcode were separated into individual BAMs respectively. Reads that failed the CCS correction were separated from those that were successfully corrected.

### Identification of insertions and insertion-mediated rearrangements after experimentally induced L1 expression

#### Shotgun whole-genome sequencing analysis

Shotgun sequencing reads were processed by the same workflow as described previously (8, 9), but with alignment (using bwa mem) to a customized reference consisting of both the primary sequences of GRCh38 and either transgene reference (Tet-On L1 or L1 GFP reporter). Haplotype-specific DNA copy-number calculation, identification of rearrangement junctions, and detection of short sequence changes (substitutions and insertion/deletion) were performed using the same workflow as described previously (8, 9).

#### Detection of de novo L1 insertions from shotgun reads

Because the L1 transgene uses a codon-optimized L1 sequence (ORFeus) that is different from endogenous L1 sequences, insertions of L1-ORFeus can be determined directly from uniquely aligned reads. De novo L1 insertions were detected using three independent methods.

First, the junctions between inserted L1s and flanking genomic DNA were identified as part of the rearrangement junction analysis (the transgene being treated as a separate contig).

Second, insertion junctions were identified from reads aligned to the L1 transgene (either Tet-On L1 ORFeus or L1 ORFeus GFP) with split subsequences or discordant pairmates aligned to the human genome. Candidate junctions were considered when there were two or more reads mapped to the L1 transgene whose pairmates or soft-clipped subsequences were mapped to genomic loci within 1kb. Candidate insertion sites with two junctions flanking an insertion (i.e., with breakpoints on opposite sides) were then manually reviewed for the presence of poly-A/T sequences at each junction. Unpaired candidate junctions were used to identify insertion-mediated rearrangements (to be described later). For RPE-1 cells with genomically integrated Tet-On L1 ORFeus, the genomic integration sites were identified by a similar approach and further validated by long reads and long-read assembly.

Third, we used xTea (10) to identify candidate sites of L1 or pseudogene insertion from reads aligned to the human genome that had discordant and soft-clipped subsequences. A candidate L1 insertion site needed to meet two criteria: 1) at least one soft-clipped or discordant read was mapped to the L1 transgene; 2) at least 60% of all soft-clipped reads were mapped to L1. Candidate sites passing these two filters were then assessed for either an insertion outcome, when a pair of sites were found within 200 bps and clipped on opposite sides, or a rearrangement junction with an L1 insertion, when unpaired. Both candidate sites of insertions (with paired junctions) or insertion-mediated rearrangements (single junctions) were further filtered by their proximity to endogenous L1 sequences. Candidate sites were removed if they met one of the following conditions: 1) each of two paired breakpoints was within 20bps of an endogenous L1; 2) both breakpoints were located in regions marked as “Simple_repeat”, “Low_complexity”, or “Other” in by repeatmasker; 3) either breakpoint of a candidate insertion site was located within 10bps of a region marked as “Simple_repeat”, “Low_complexity” or “Other” by repeatmasker.

In each of the three analyses, candidate insertion junctions and singleton junctions were detected from individual samples (including control samples), but the supporting reads were collected from all samples. Junctions identified in more than one sample or having supporting reads from more than one sample (indicating that they were either ancestral alterations or recurrent technical artifacts) were excluded.

#### Detection of de novo pseudogene insertions from shotgun reads

Insertions of processed pseudogenes were also detected using three independent approaches.

First, insertions were identified as part of the rearrangement detection workflow from both discordant/split reads spanning exonic junctions at the source gene locus and from discordant/split reads at the insertion sites.

Second, SideRetro(11) was used to identify candidate insertion sites using the parameter ‘mc-m3-x200k’.

Third, xTea (10) was run to identify candidate sites of pseudogene insertions from reads aligned to the human genome that have discordant and soft-clipped subsequences. Candidate pseudogene insertion junctions were selected based on the following criteria: 1) there were two adjacent breakpoints (within 200bps) consistent with an insertion; 2) neither breakpoint resided within repeats (same as above for detecting L1 insertions); 3) no other breakpoint was detected within 10bps from the candidate breakpoints; 4) the soft-clipped sequences and the discordant pairmates of supporting reads at each breakpoint must have aligned to locations within 20kb in the human genome; 5) the mapping locations of clipped parts of supporting clipped reads at two ends of a candidate insertion were within 5kb and the mapping locations of mates of discordant supporting reads at two ends of a candidate insertion were within 10kb.

Candidate sites output by SideRetro (11) and using xTea (10) with filtering were intersected to generate the list of candidate insertions, and the list of candidate insertions was merged with the list of junctions revealed from the rearrangement analysis. After excluding insertions with read support from more than one sample, the remaining insertions were manually reviewed and curated with help from long-read data and RNA-Seq data.

#### Processing of PacBio long-read data

Error-corrected (Hi-Fi) reads were aligned to the same references (GRCh38 + either L1 transgene) using minimap2 (version 2.26-r1175) with the following parameters “-a -k19 -w19 - U50,500 -g10k -A1 -B4 -O6,26 -E2,1 -s40 -Y -c.” The only difference from the preset “--map-hifi” as suggested for the alignment of PacBio Hifi reads was ‘-s40’ instead of ‘-s200.’ The addition of ‘-c’ and ‘-Y’ was used for downstream processing of aligned reads.

We performed haplotype-resolved (diploid) assembly of long reads from three Dox clones (with Tet-On L1 ORFeus) and previously published Hi-C data using hifiasm (8, 9). We used minimap2 (-asm5) to align the assembled contigs to the Tet-On L1 transgene to identify contigs with genomically integrated Tet-On transgenes, and then determine the GRCh38 coordinates of the insertion sites by aligning these contigs to the human genome reference.

#### Manual curation of insertions and insertion-mediated rearrangement junctions

We employed the following criteria/strategies to validate/refine L1/pseudogene insertions or insertion-mediated rearrangements identified from short reads: (1) The insertion was supported by at least one long read when long-read data were available (see e.g., **Figure 3A**). (2) When long-read data were unavailable, two breakpoints need to be identified at the target site, including one with soft-clipped poly-A/T sequences (the 3’-junction of the insertion); moreover, no copy-number changes were permitted at the breakpoints (based on 90kb-level haplotype-specific DNA copy number data as well as local sequence coverage). (3) For pseudogene insertions, reads derived from the inserted sequence (both short and long reads) were often misaligned to endogenous retrocopies of the same source gene. To resolve such ambiguity, we looked for the source gene and verified the 5’ and 3’ junctions from the short read data. We further validated expression of the source gene using the RNA-Seq data. (4) For L1-mediated reciprocal translocations, the features were similar to (2) except that the two junctions had different translocation partners. (5) For L1-mediated rearrangement junctions with unbalanced translocations, the breakpoint orientation (+) or (-) needed to be consistent with the directionality of copy-number change as assessed from the sequence coverage. (6) All copy-number changepoints between segments of 1Mb or higher were manually reviewed to identify additional breakpoints that were missed by the automatic rearrangement and insertion detection method.

Identification of 5’-inverted insertions was based on the orientation of the split/discordant subsequences to the L1 transgene reference and the split alignments of the insertion sequence resolved by long reads. The internal junction of a 5’-inverted insertion was either resolved by long reads or identified based on proximity to the breakpoint of the twin-primed reverse transcription (see **Fig.2C**).

### Analysis of insertions and insertion-mediaged rearrangements

#### Microhomology and untemplated insertions at junctions

Microhomology or untemplated insertions at any rearrangement junction (including insertion junctions) were calculated based on the lengths of aligned subsequences of soft-clipped reads. Microhomology/untemplated insertions at the 3’-end (poly-A) of retrocopied sequences were not assessed.

#### Calculation of insertion lengths and target site deletion/duplication sizes

For L1 insertions without 5’ inversions, the insertion length was calculated from the 5’ and 3’-breakpoints in the L1 reference (excluding the poly-A sequence). For 5’-inverted L1 insertions, the insertion length was calculated as the sum of two inverted insertions. For pseudogene insertions, the insertion length was calculated from the number of bases from the inserted sequence (resolved by long reads) that were aligned to the source gene locus. When long-read data were unavailable, the insertion length was calculated similarly as for L1 insertions except that the introns were excluded.

The length of target site duplication or deletion was calculated (1) from the breakpoints at the target site, or (2) from the insertion sequence resolved by long reads.

---

**Figure S1.**
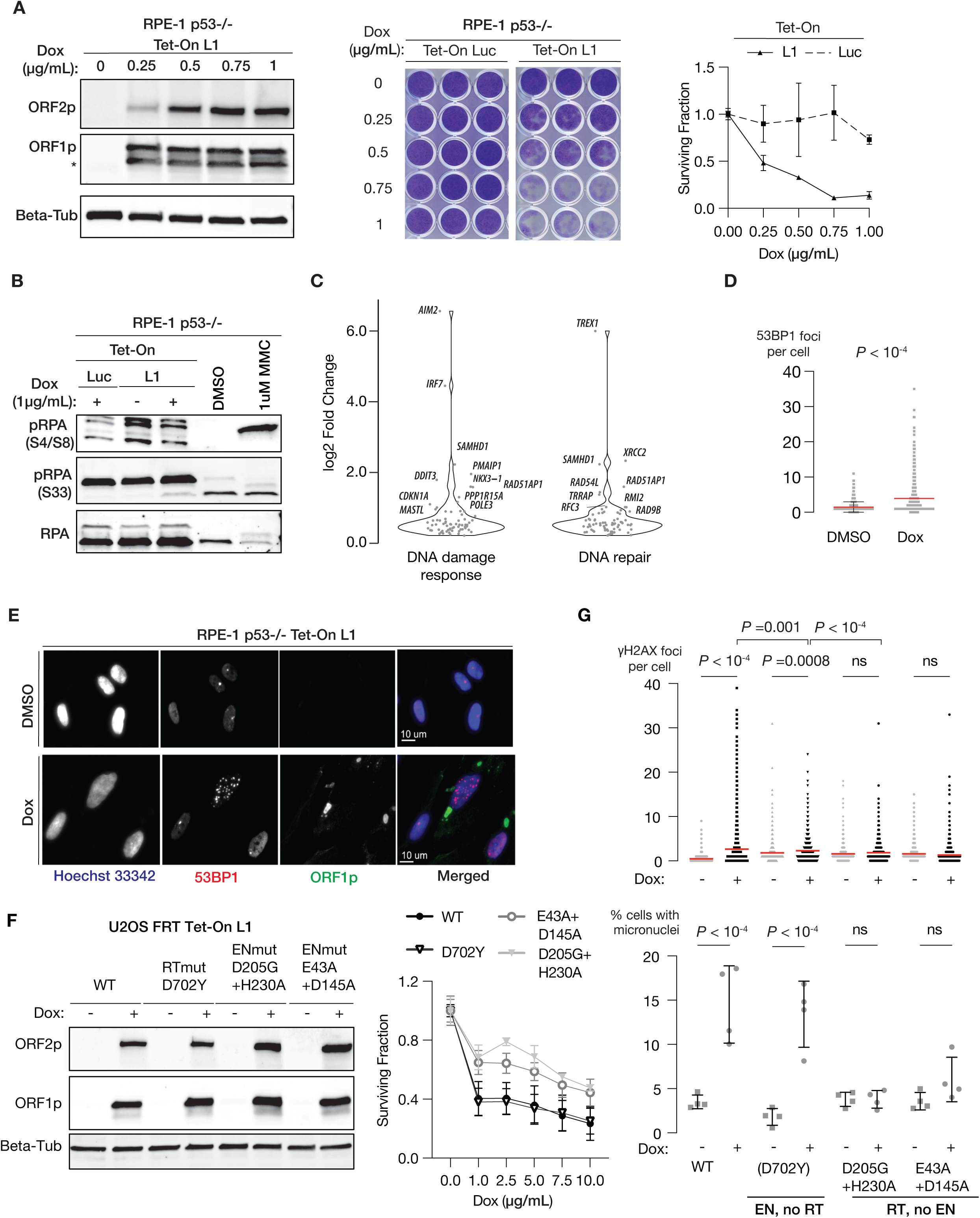
Further evidence of DNA damage and DSB generation from L1 expression. **A.** Left: Immunoblot of ORF1p and ORF2p expression in p53-null RPE-1 cells with Tet-On L1 after treatment with Doxycycline at different concentrations. Middle: Reduced colony formation in cells with Dox-induced L1 expression. Right: Quantification of the relative abundance of cells after L1 induction in comparison to the control (Dox-induced luciferase expression). **B.** Immunoblots of pRPA (S4/S8) and pRPA (S33) from whole cell lysates after L1 induction and from control experiments, similar to **Figure 2B**. **C.** Log2 fold changes of upregulated genes in the DNA damage response and DNA repair pathways (based on Gene-Ontology). Related to **Figure 2C**. **D.** Increased 53BP1 foci in p53-null RPE-1 cells after L1 induction (612 cells under Doxycycline) relative to control cells (538 cells with DMSO treatment). Two independent experiments; *P* <0.0001; two-tailed Mann-Whitney U-test. **E.** Representative immunofluorescence images of 53BP1, ORF1p, and Hoechst 33342 in p53-null RPE-1 with (Dox) or without (DMSO) L1 induction. Bar scale: 10µm. **F.** Validation of L1 induction in U2OS cells with either wild-type or mutant L1 with a Tet-On promoter integrated at an FRT locus. Left: Immunoblots similar to **A**. Right: Reduced cell survival under the induction of wildtype or EN-proficient/RT-mutant L1 in comparison to the induction of L1 with inactivated EN (*n* = 6 replicates except for ENmut, D205G:H230A). **G.** Quantification of γH2AX foci (top) and micronucleation frequency (bottom) in U2OS FRT cells with Dox-induced expression (1 µg/mL) of wildtype and mutant L1. Two independent experiments for each condition. Induction of wildtype L1 (first group) produces the most significant increase in γH2AX foci and micronucleation. Induction of L1 with proficient EN but inactive RT (second group) produces an attenuated but significant increase in γH2AX foci and in micronucleation frequency. By contrast, induction of L1 with proficient RT but inactive EN produces no noticeable change in either γH2AX or micronucleation relative to control. P-values are calculated using one-way ANOVA with Tukey test.

**Figure S2.**
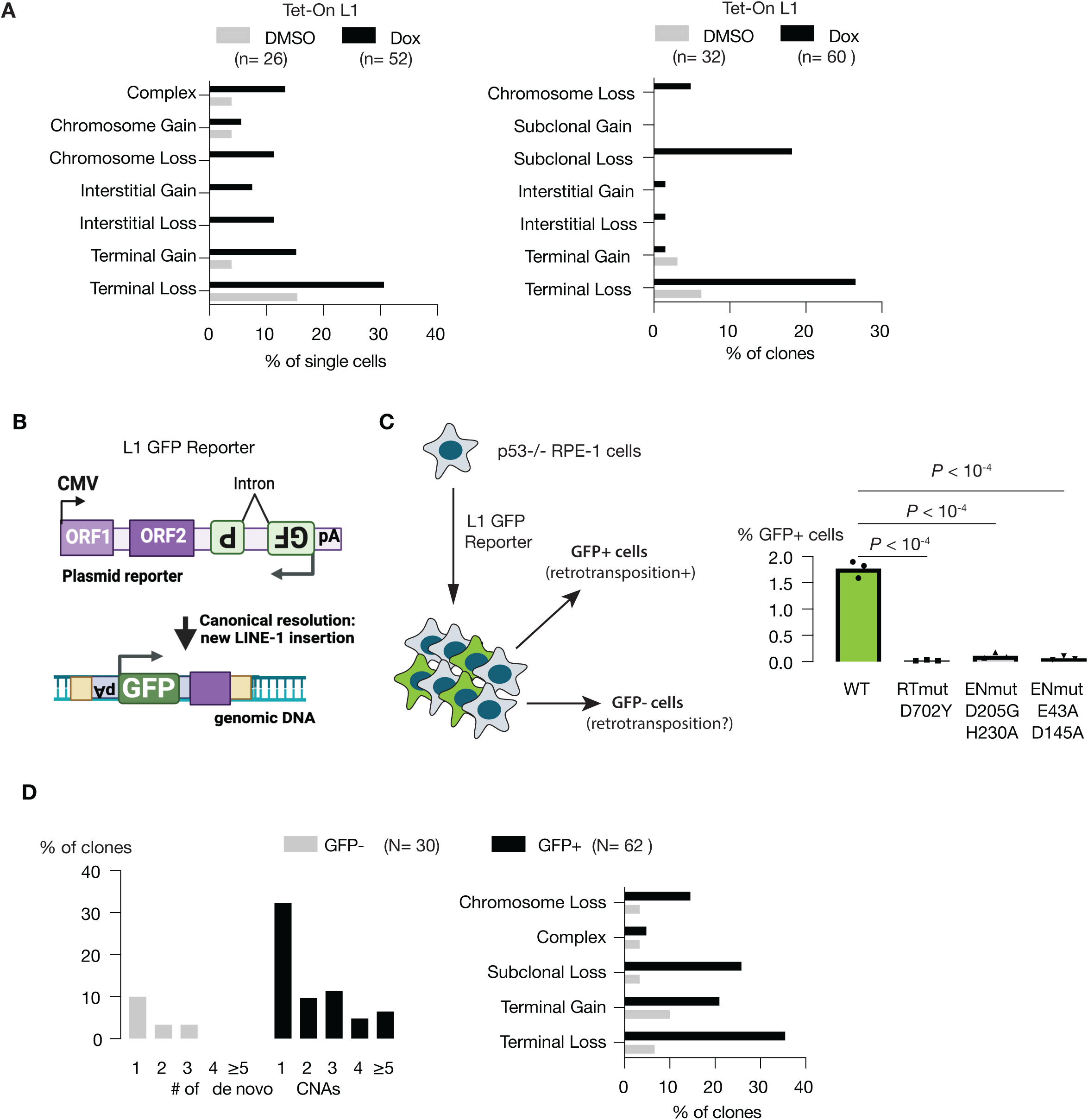
Large segmental copy-number alterations in single cells or progeny derived from single cells after L1 expression. **A.** Large segmental CNAs in single cells (left) and single-cell derived clones (right) after Doxycycline-induced L1 expression. Only alterations of segments ≥ 5Mb are counted; CNAs on different parental chromosomes are evaluated separately. **B.** Schematic diagram of the L1 GFP reporter. The anti-sense split GFP gene in the 3’-UTR can only be transcribed from the retrocopy generated by retrotransposition. Therefore, GFP+ cells must have had one or multiple integrations of the split GFP gene. GFP-cells may also contain truncated copies of the reporter. **C.** Frequency of L1 retrotransposition in p53-null RPE-1 cells assessed using the L1 GFP reporter. Inactivation of reverse transcriptase activity completely abolishes retrotransposition, whereas inactivation of endonuclease activity suppresses but does not eliminate retrotransposition. Three replicates in each condition. *P*-values are calculated using one-way ANOVA with Tukey test. **D.** Large segmental CNAs in clones expanded from single GFP- and GFP+ cells. Similar to **A**. See **Supplementary Figure 1** for a summary of large CNAs on each chromosome in each sample.

**Figure S3.**
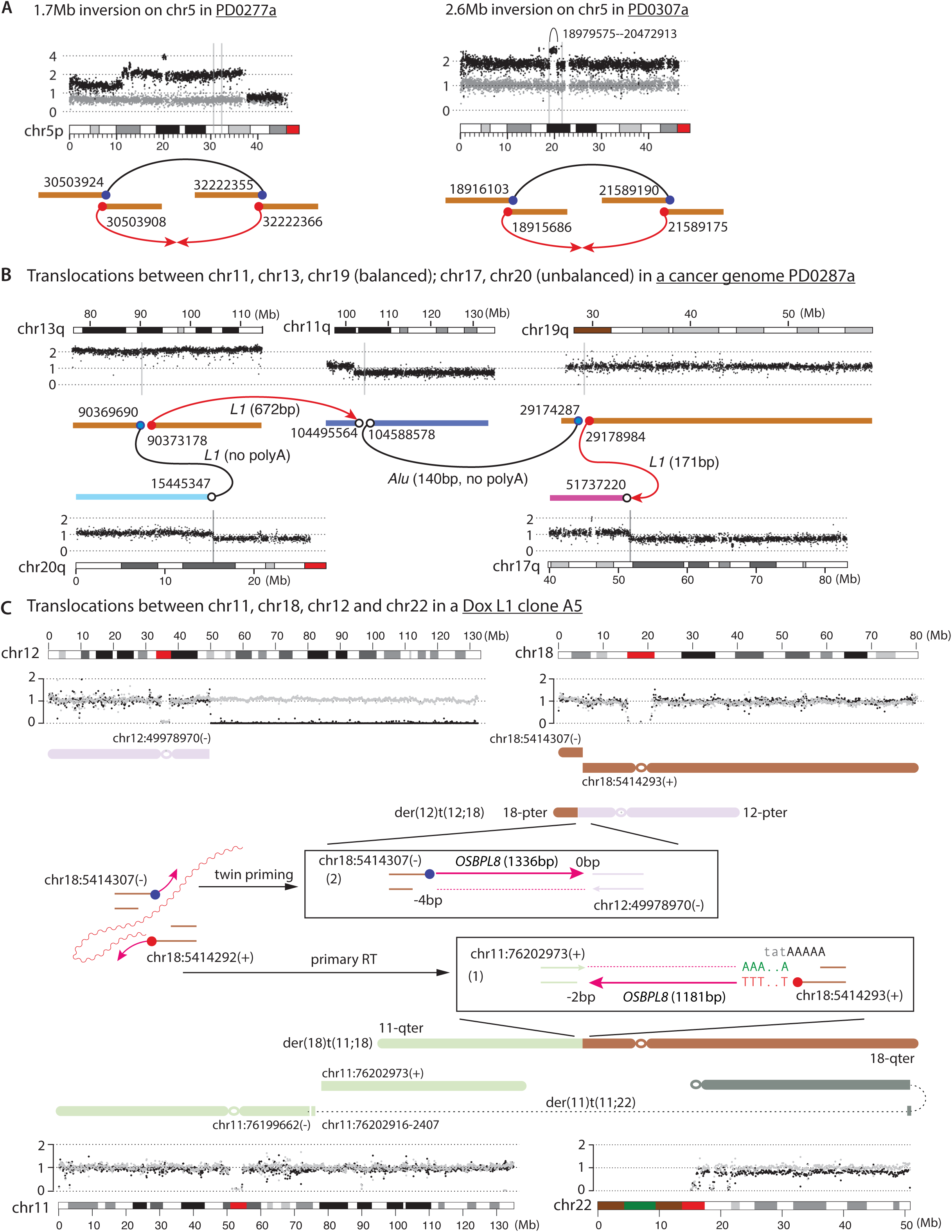
Large inversions and chromoplexy from ORF2p-induced DSBs. **A.** Two instances of large inversions identified in cancer genomes reflecting a similar DSB exchange outcome as reciprocal translocations. In both instances, L1 insertions from both DSB sites (red arrows) are retained in a single inversion junction. The DNA copy number show no transition on either homolog at both sites (vertical gray lines). In the instance from PD0307a, the inversion breakpoints near 18.91Mb are close to a tandem duplication (18.98-20.47Mb) that is likely unrelated to the inversion. **B.** Four translocations between three pairs of reciprocal breakpoints (on chr11, chr13, and chr19) and two unbalanced breakpoints (on chr20 and chr17) in a cancer genome (PD0287a). Polyadenylated L1 insertions are present in two junctions (red arrows) and identify two primary RT breakpoints (red filled circles) on chr13 and chr19; the other two junctions contain non-polyA insertions (black). The copy-number plots for chr20 and chr17 show only the translocated homologs; for chr13, the other homolog is deleted; for chr11, both homologs show the same copy number (only one is shown); for chr19, the other homolog is amplified (not shown). On chr13 and chr19, the gaps between the reciprocal breakpoints (3.5kb on chr13 and 4.7kb on chr19) are most likely due to resection of the reciprocal end (light blue circles) indicating that these junctions are formed in S/G2. There is a larger deletion between the reciprocal breakpoints on chr11 (93kb). **C.** Three translocations between chr11, chr18, chr12, and chr22 in a Dox L1 clone (A5). Copy-number data are similar to **Figure 3**. Two cDNA sequences derived from the *OSBPL8* mRNA (chr12:76,351,797-76,352,977 and chr12:76,352,990-76,354,325) are found at both breakpoints on chr18: The polyadenylated insertion identifies the primary RT breakpoint (5,414,292); the insertion at the other breakpoint is consistent with twin priming. Had these two breakpoints been ligated together, they would have generated a 5’-inverted L1 insertion for which the twin priming model was originally proposed to explain. This example thus provides a snapshot of an intermediate step of the twin-priming model. For the two breakpoints on chr11, the breakpoint at 76,199,662(-) is next to an inversion of chr11:76,202,407-76,202,916, which likely arises from DSB end remodeling (see **Figure 4**). The centric chr11 segment is ligated to chr22q that has a terminal deletion flanked by an inverted duplication, forming a dicentric chromosome; there is subclonal copy-number loss between the two centromeres (black dots) reflective of DNA loss from the chromosome-type BFB cycles (also see **Figure 7**).

**Figure S4.**
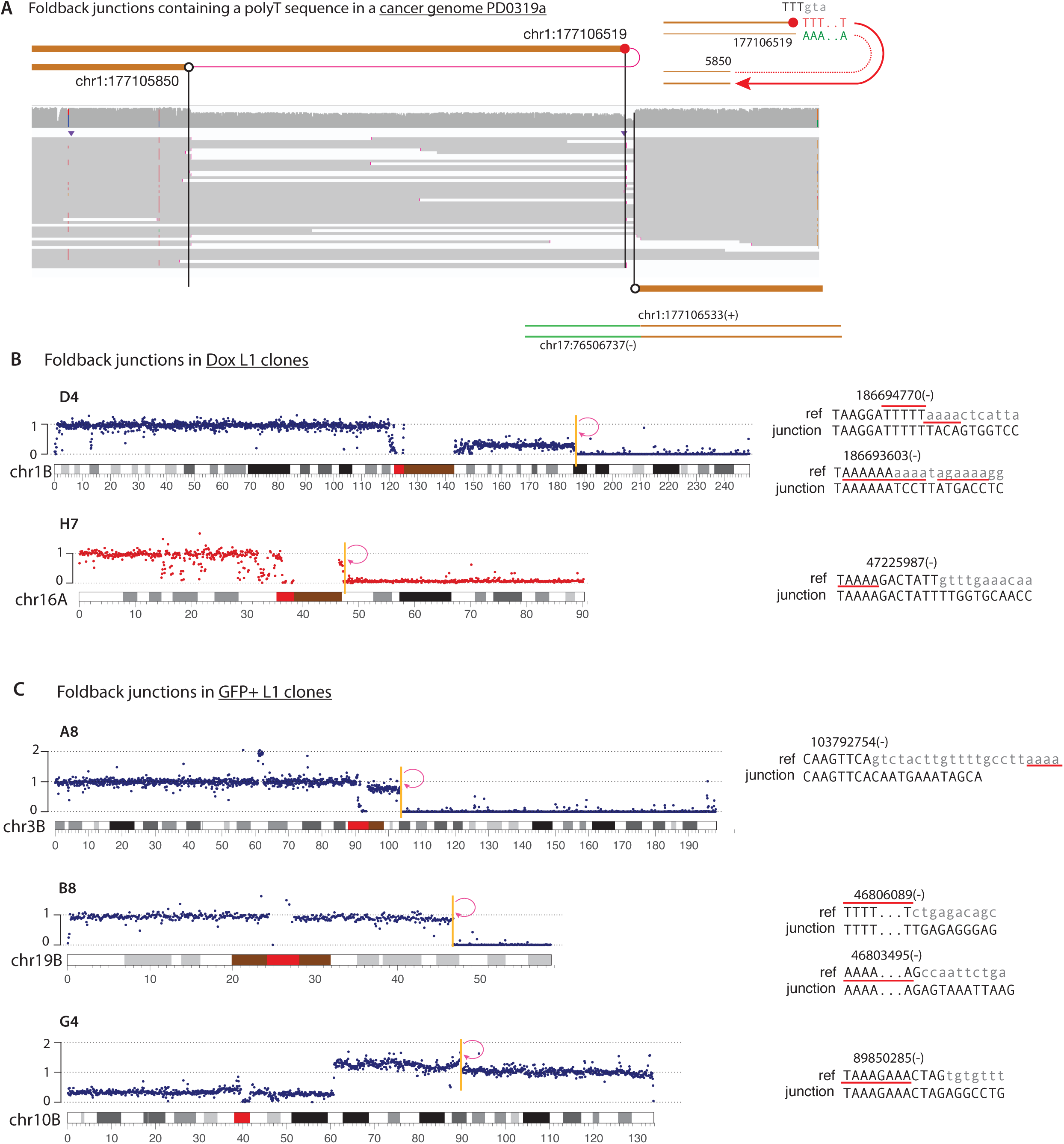
Additional examples of foldback junctions in cancer genomes and experimental L1 clones. **A.** Another example of foldback junction with a reciprocal breakpoint identified in a cancer genome. Here the reciprocal breakpoint forms a simple translocation junction. **B.** Two examples of foldback junctions identified in Dox L1 clones. **C.** Three examples of foldback junctions identified in GFP+ L1 clones. Although the foldback junctions shown in **B** and **C** do not contain TPRT insertions, the presence of ORF2p EN target sites near the breakpoints suggests that the foldbacks can arise from the reciprocal end generated by ORF2p.

**Figure S5.**
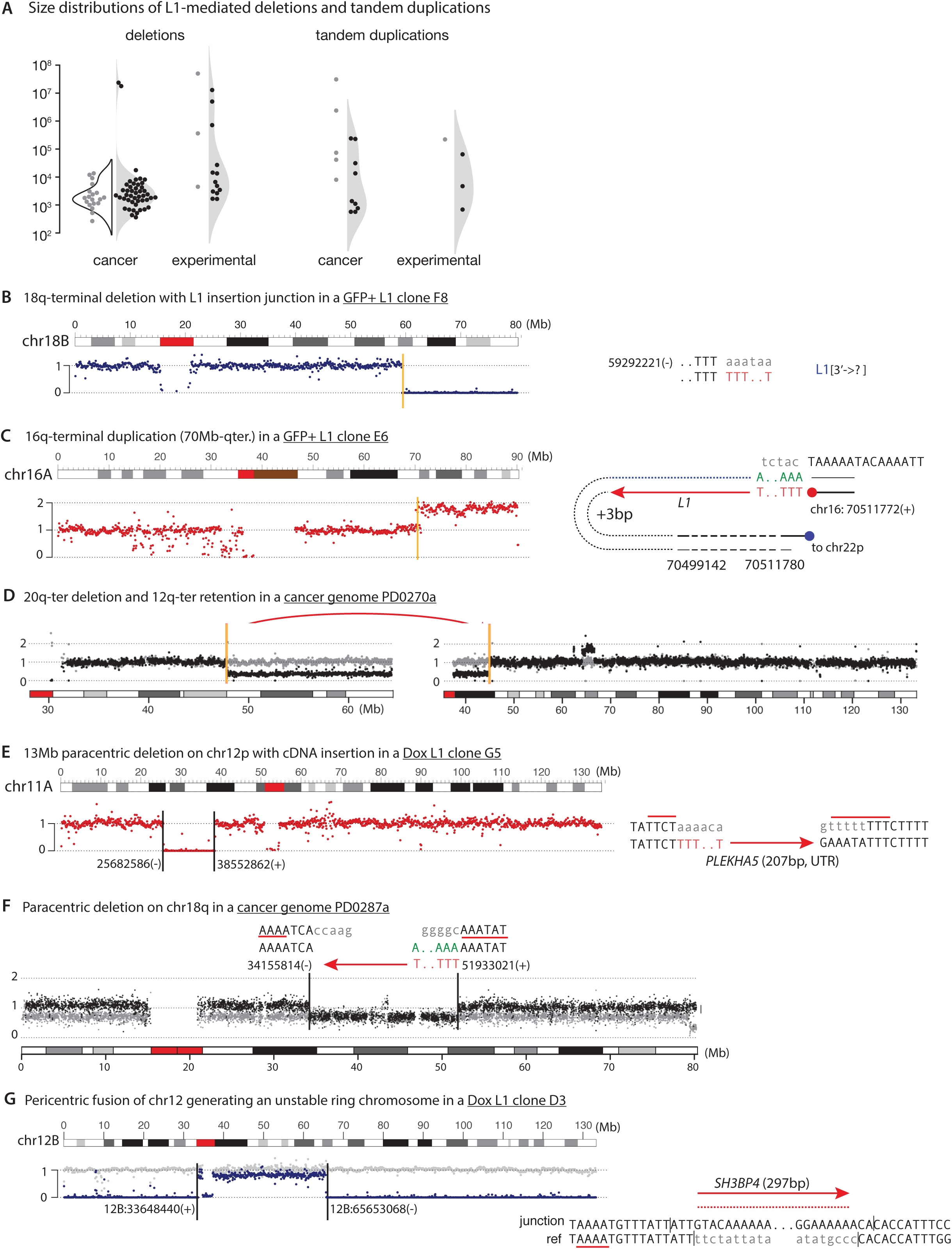
Simple segmental copy-number alterations arising from L1-mediated translocations or rearrangements. **A.** Size distributions of simple deletions and tandem duplications with rearrangement junctions containing L1 insertions or displaying TPRT features. For the instances from cancer genomes, black dots represent junctions containing polyadenylated L1 insertions; gray dots represent junctions containing non-polyadenylated insertions. For the instances from experimental L1 clones (all from Dox-L1 clones), black dots represent junctions containing polyadenylated L1 insertions or with one or both breakpoints adjacent to ORF2p target sequences; gray dots represent the remaining simple deletions/tandem duplications. Observations from both cancer genomes and experimental L1 clones suggest short deletions (1-10kb) as the most frequent outcome of L1-induced DSB. **B** and **C**. Examples of terminal deletion (**B**) and duplication (**C**) identified in experimental L1 clones that we infer to arise from a secondary deletion or duplication of stable chromosomes generated by L1-mediated translocations. Shown are the copy number data of the translocated homologs only. Note the constant copy number of the retained, deleted, or duplicated segments as expected for stable translocations. In **C**, two breakpoints are detected near the duplication boundary: the primary RT end gives rise to the breakpoint at chr16:70,511,772(+), whereas the reciprocal end gives rise to the translocation breakpoint chr16:70,511,780(-). The 12kb segment 70,499,142-70,511,780 could have resulted from cleavage of a large 3’-overhang. **D**. Two terminal deletions inferred to reflect a chromosome deletion after L1-mediated translocations in a cancer genome. Black and gray dots represent haplotype-specific DNA copy number and the red arrow represents the L1 insertion linking the 20q and 12q segments. **E** and **F.** Large paracentric deletions identified in an experimental L1 clone (**E**) and in a cancer genome (**F**). In **E**, a 13Mb internal deletion on 12p contains insertion of a truncated *PLEKHA5* cDNA between the breakpoints. Both breakpoints are adjacent to ORF2p EN target sequences, supporting an origin from ORF2p-induced DSB. **F** is similar to **E**, except that the first cDNA strand is on the reverse strand. **G**. A ring chromosome inferred to be present in an experimental L1 clone. The two terminal breakpoints are joined together with insertion of a *SH3BP* cDNA, which would result in a ring chromosome. The inference of a ring chromosome is supported by the subclonal loss of this chromosome (relative to the intact homolog shown in gray).

**Figure S6.**
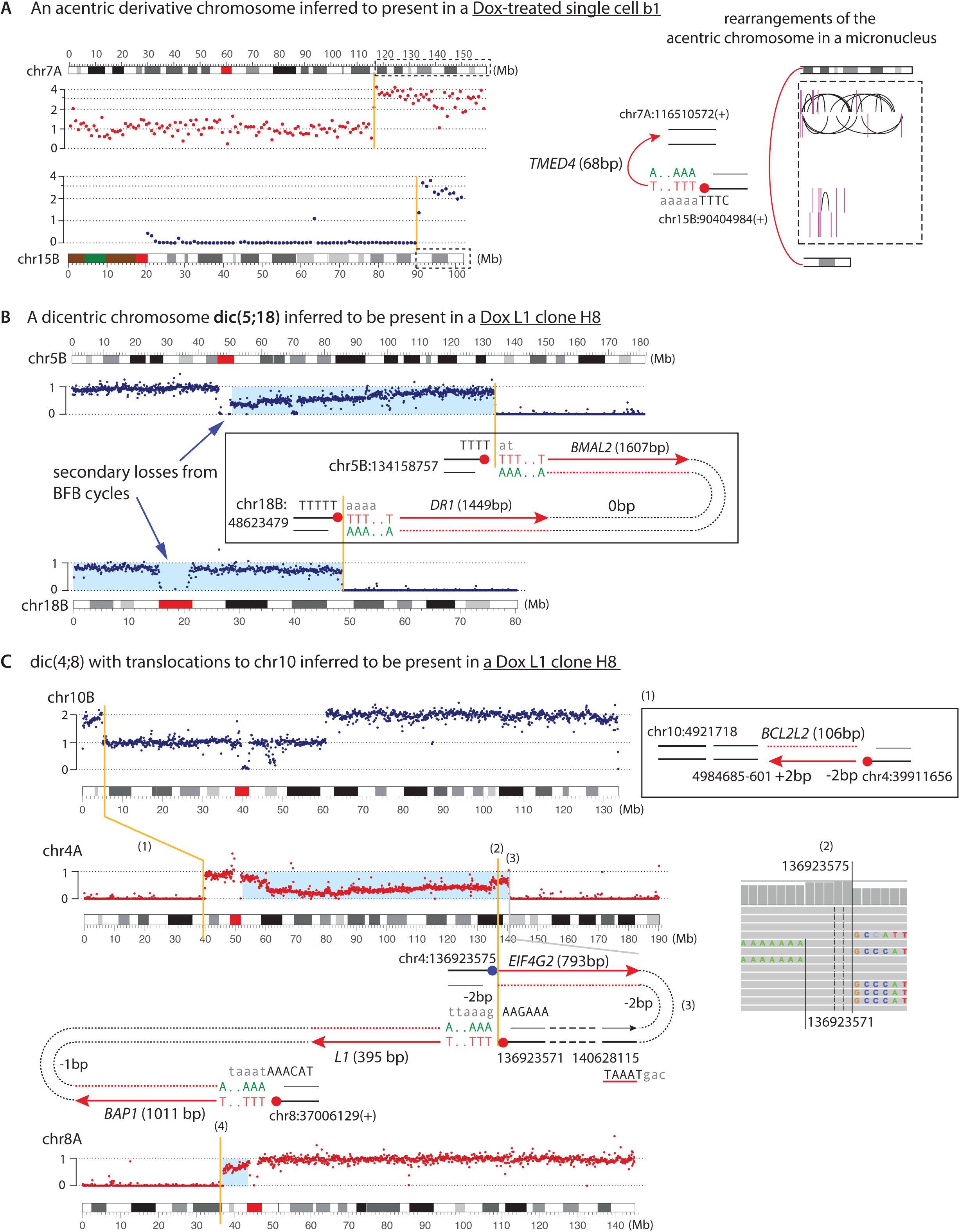
Examples of chromosomal instability after L1-mediated translocations identified in single cells or clones with experimentally induced L1 expression. **A.** An example of acentric chromosome inferred to have been generated by L1-mediated translocation in a single Dox L1 cell b1. The inference of an ancestral acentric chromosome is based on (1) the structure of the segmental gain (on 7q) and retention (on 15q) based on haplotype-specific DNA copy number (7A and 15B); (2) the presence of intra- and inter-chromosomal rearrangements restricted to these two regions, indicating chromothripsis of an acentric chromosome partitioned in a micronucleus. A truncated poly-adenylated cDNA copy of the *TMED4* gene identifies the ancestral primary RT end to be chr15:90,404,984. **B.** A dicentric chromosome generated by L1-mediated translocation inferred to be present in the Dox L1 clone H8. Shown are the haplotype-specific DNA copy number of the translocated homolog (5B and 18B). The inference of dic(5;18) is based on three observations: (1) a single breakpoint on each arm of the translocated chromosome; (2) deletion of sequences telomeric to the breakpoints; and (3) subclonal copy-number loss on the dicentric chromosome (highlighted in blue), including segmental losses between the two centromeres, as expected for the chromosome-type BFB cycles. Both breakpoints are inferred to have derived from the primary RT ends (red filled circles) based on TPRT signatures. **C.** Another dicentric chromosome inferred to be present in the Dox L1 clone H8. This dicentric chromosome contains two inter-chromosomal translocation junctions. The junction between the 4q breakpoint [chr4:136923571(+)] and the 8p breakpoint [chr8:37006129(+)] contains two independent TPRT insertions that establish their origin from primary RT ends generated by ORF2p. The reciprocal breakpoint chr4:136923575(-) (see IGV screenshot showing a 5bp target-site duplication) most likely derives from the reciprocal end and is extended by reverse transcription using a different mRNA (*EIF4G2*). This scenario is similar to twin priming (**Figure S3C**) except that the two ends are extended using different mRNA templates and ligated to distal DNA ends. Finally, the terminal 4q breakpoint [chr4:140628115(-)] is adjacent to an ORF2p EN target sequence (T|AAA) and could have derived from the reciprocal end generated by ORF2p. The 4p breakpoint [chr4:39911656(+)] is extended by reverse transcription of the *BCL2L2* mRNA from within the 3’-UTR (no poly-A) but is not adjacent to ORF2p EN target sequences. The inference of dic(4;8) is based on subclonal segmental copy-number losses between the two centromeres and the complete deletion of terminal sequences.

**Figure S7.**
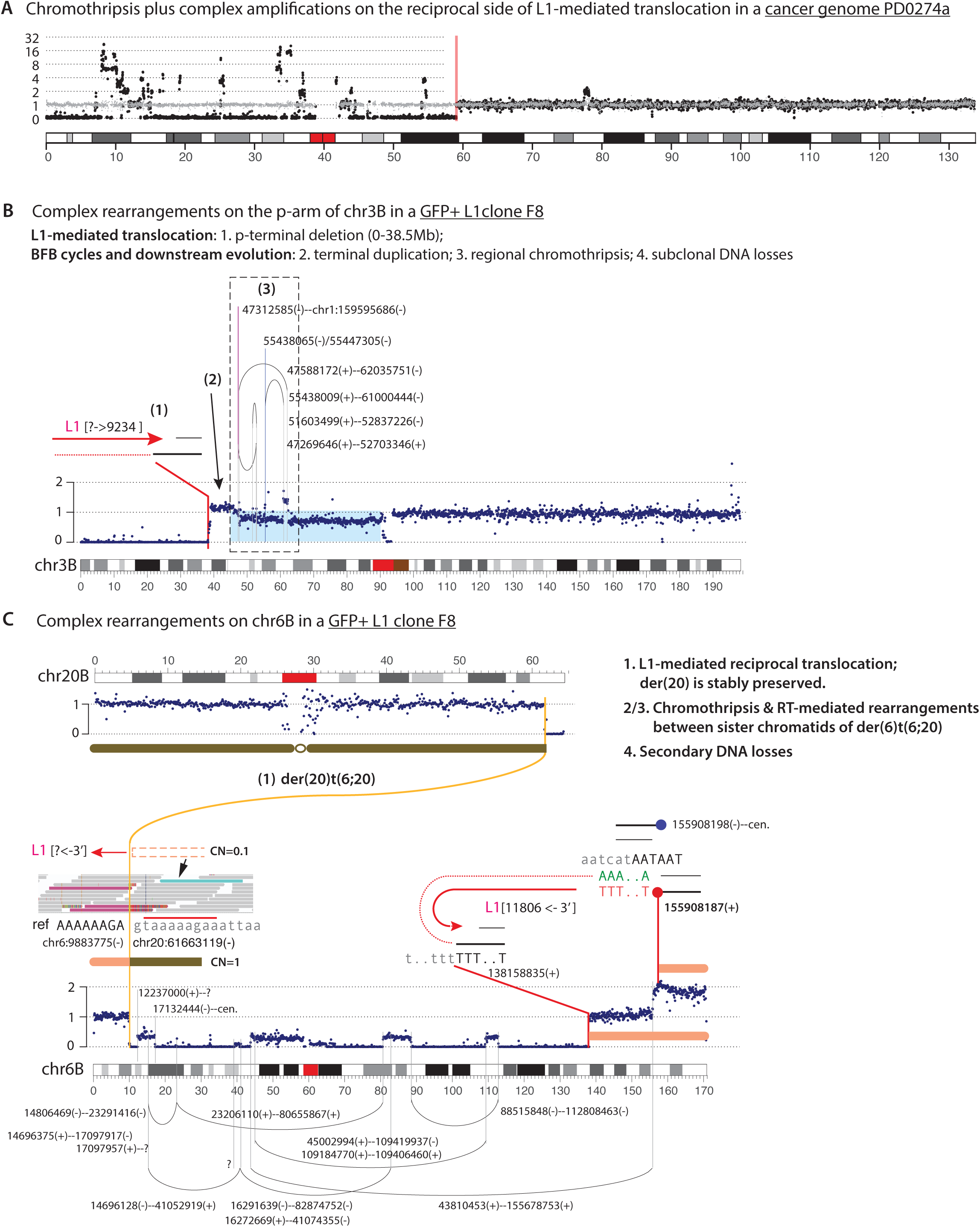
Examples of chromothripsis and complex rearrangements arising downstream of L1-mediated translocations. **A.** An example of chromothripsis with complex amplifications in a cancer genome. Black and gray dots represent the haplotype-specific DNA copy number (10kb bins). The black homolog is inferred to be the homolog that underwent a L1-mediated translocation (breakpoint shown as the vertical red line). Note the complex SCNA pattern to the left (i.e., on the reciprocal side) of the breakpoint. **B.** An example of chromothripsis on chr3p identified in the GFP+ L1 clone F8. A L1-mediated translocation (the translocation partner cannot be determined) leads to the p-terminal deletion, followed by one or multiple BFB cycles, creating subclonal DNA losses and complex rearrangements on the 3p arm. The L1-mediated translocation is inferred to be the initiating event of complex rearrangements. **C.** An example of chromothripsis of chr6 in the same L1 clone as in **B**. The first event is a translocation between chr6 and chr20. We infer there to be a stable der(20)t(6;20) between the 6p terminal segment (chr6:0-9,883,775). Although the translocation junction does not contain any insertion, there is a single read to the right of the chr6 breakpoint (shown in light green) indicating an L1-mediated translocation junction as well as partial retention of sequences to the right of the chr6 breakpoint (with clonal fraction ∼10%). These observations support an origin of the chr6:chr20 translocation from an ORF2p-induced DSB on chr6. We also observe an L1-mediated junction between two breakpoints on 6q [138,158,835(+) and 155,908,187(+)]. The L1 insertion identifies chr6:155,908,187 as the primary RT breakpoint that has a reciprocal breakpoint at 155,908,198(-); the two breakpoints display a 12bp TSD feature. Based on the duplication of the q-terminal segments, we infer the two breakpoints to originate from DNA ends on sister chromatids. We note that the copy-number transition occurs at the breakpoint 155,678,753(+).

**Supplementary Figure 1.**
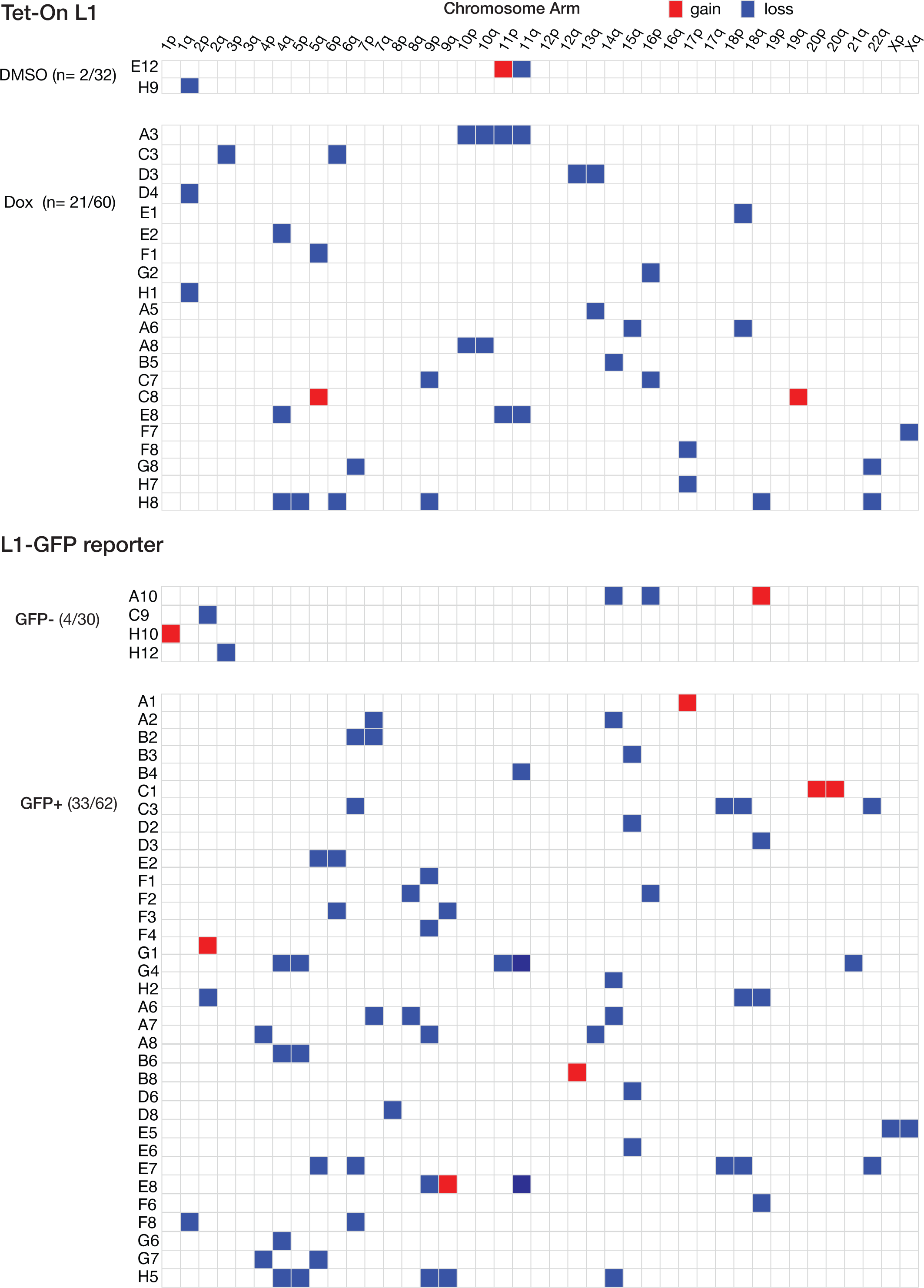
Heatmap of de novo large CNAs (>5Mb) in clones derived from single cells with induced L1 expression and from control cells. Top: control and Dox L1 clones from the Tet-On L1 system, related to **Figure S2A**. Bottom: GFP- and GFP+ clones from the L1-GFP reporter system, related to **Figure S2D**. Only clones with detectable large CNAs are shown.

**Supplementary Figure 2.**
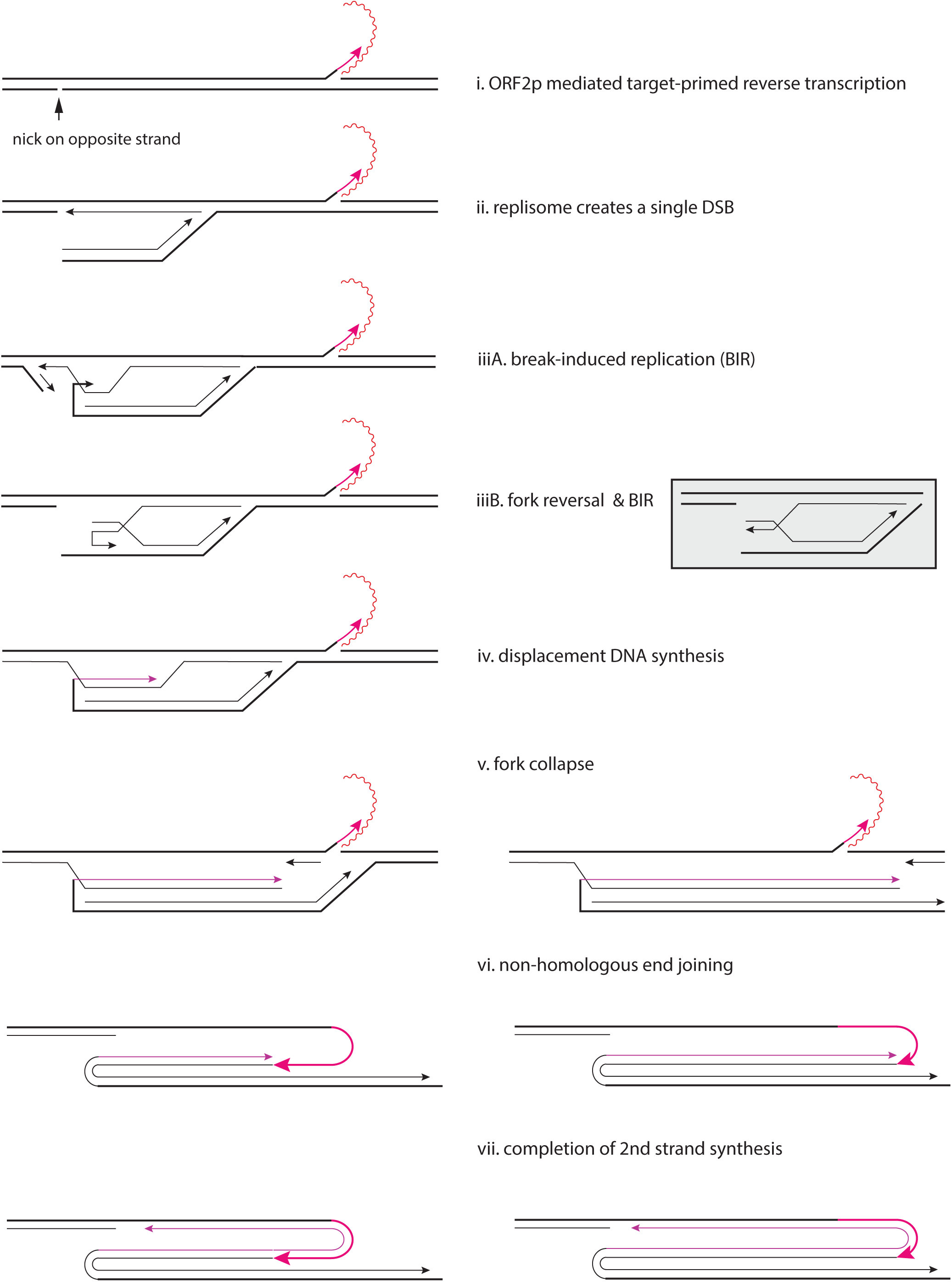
A proposed model for the generation of triplications flanked by foldbacks based on the presence of TPRT insertions. We propose that the triplication arises from re-replication (iii-v) initiated from unpaired DNA ends generated at nicks on opposite template DNA strands (i-ii); one foldback (left) arises from strand invasion (iii) that may be promoted by the presence of inverted repeats (palindromic sequences); the other foldback (right) arises from end-joining. Note that the model shown assumes that the replication fork proceeds past nicks on the lagging strand template (iii) but collapses at nicks on the leading strand template (v). The non-homologous end-joining step (vi) is proposed based on the observation that the L1-mediated foldback junctions (**Figure 5C**) are non-homologous.

**Supplementary Figure 3.**
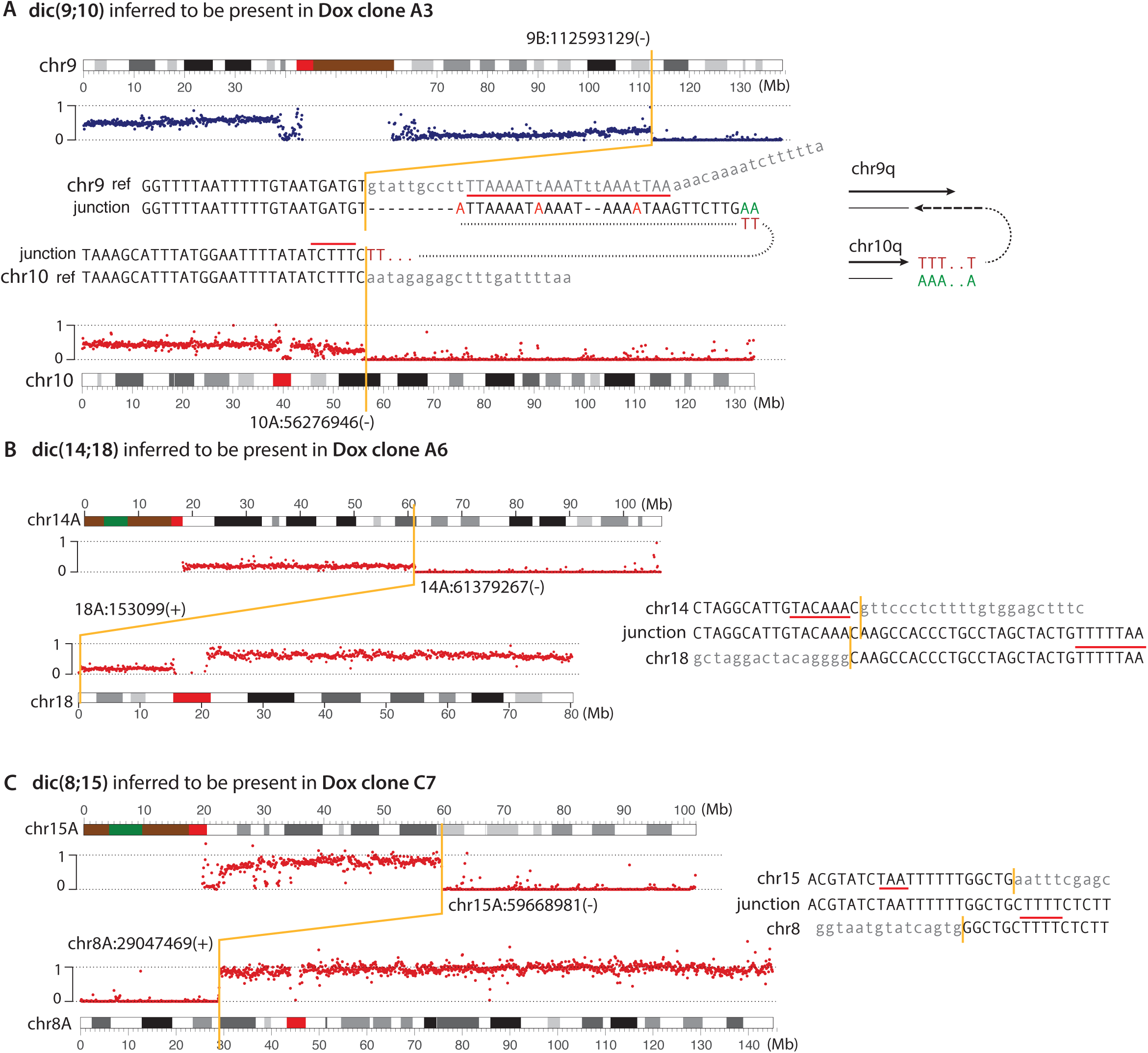
Dicentric chromosomes inferred to be present in Dox L1 clones. The copy number plots show the normalized copy number of the translocated homolog. The inference of dicentrics is based on (1) deletions of telomeric sequences on both chromosomes; (2) subclonal DNA losses between the two centromeres (the outcome of the chromosome type BFB cycles). In the example shown in **A**, the junction between chr9 and chr10 contains both a poly-A sequence and a sequence that is homeologous to the flanking sequence on the chr9 breakpoint; the poly-A sequence and the ORF2p EN target sequence near the chr10 breakpoint identifies this breakpoint’s origin from an ancestral primary RT end; the presence of ORF2p EN target sequences near the chr9 breakpoint (on opposite strand) indicates that the chr9 breakpoint likely derived from an ancestral reciprocal end. For the two examples shown in **B** and **C**, we identify possible ORF2p EN target sequences near both breakpoints that support their origin from ancestral reciprocal ends.

**Supplementary Figure 4.**
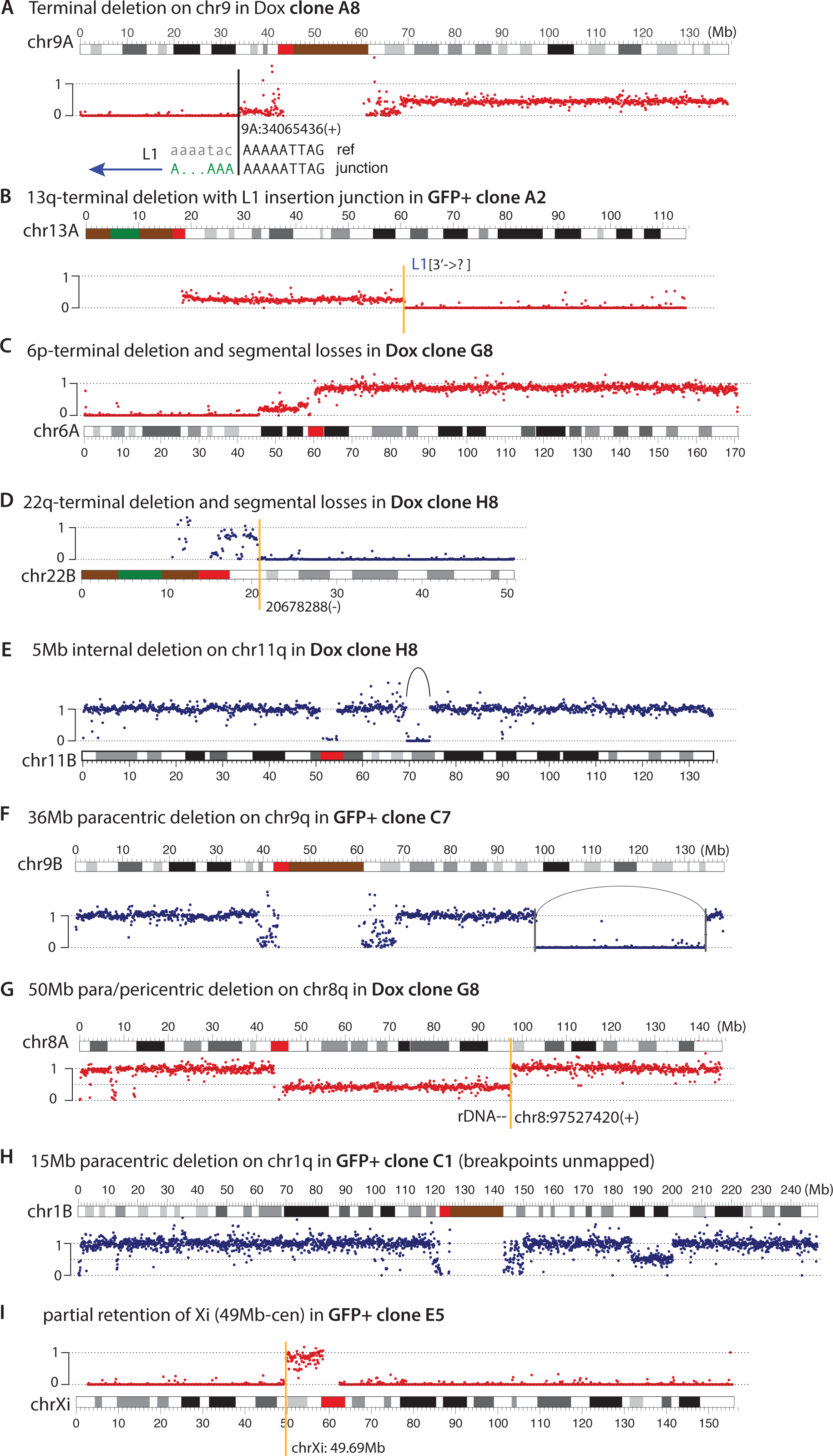
Additional examples of large segmental CNAs in experimental L1 clones. The copy number plots show only the normalized copy number of the altered homolog. In the examples shown in **A** and **B**, we identified L1 insertions at the terminal deletion boundary but could not map the partner locus of the translocation as long read data are unavailable. In the remaining samples, we did not identify L1/cDNA insertions at the CNA boundaries or were unable to map the breakpoints precisely.

**Supplementary Figure 5.**
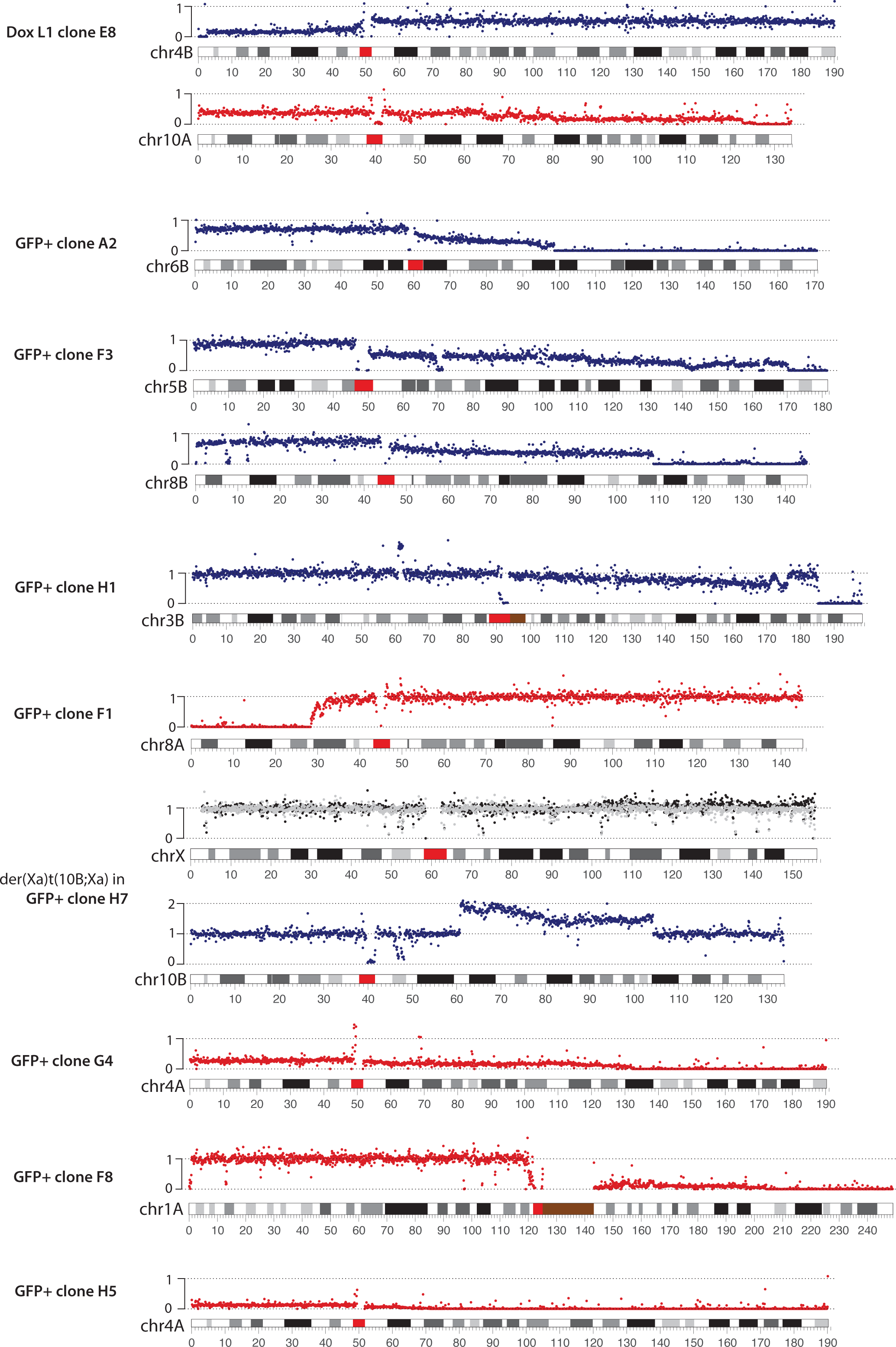
Examples of sloping copy-number variation indicating BFB cycles in experimental L1 clones. The copy number plots show only the normalized copy number of the altered homolog. For the example in GFP+ clone H7, the sloping copy-number variation is on the extra copy of the 10q segment that is appended to the active X. Note the minor copy-number gain in Xa (black dots above the gray from the normal homolog) and copy-number losses in the 10q arm relative to trisomy in the parental line.

**Supplementary Figure 6.**
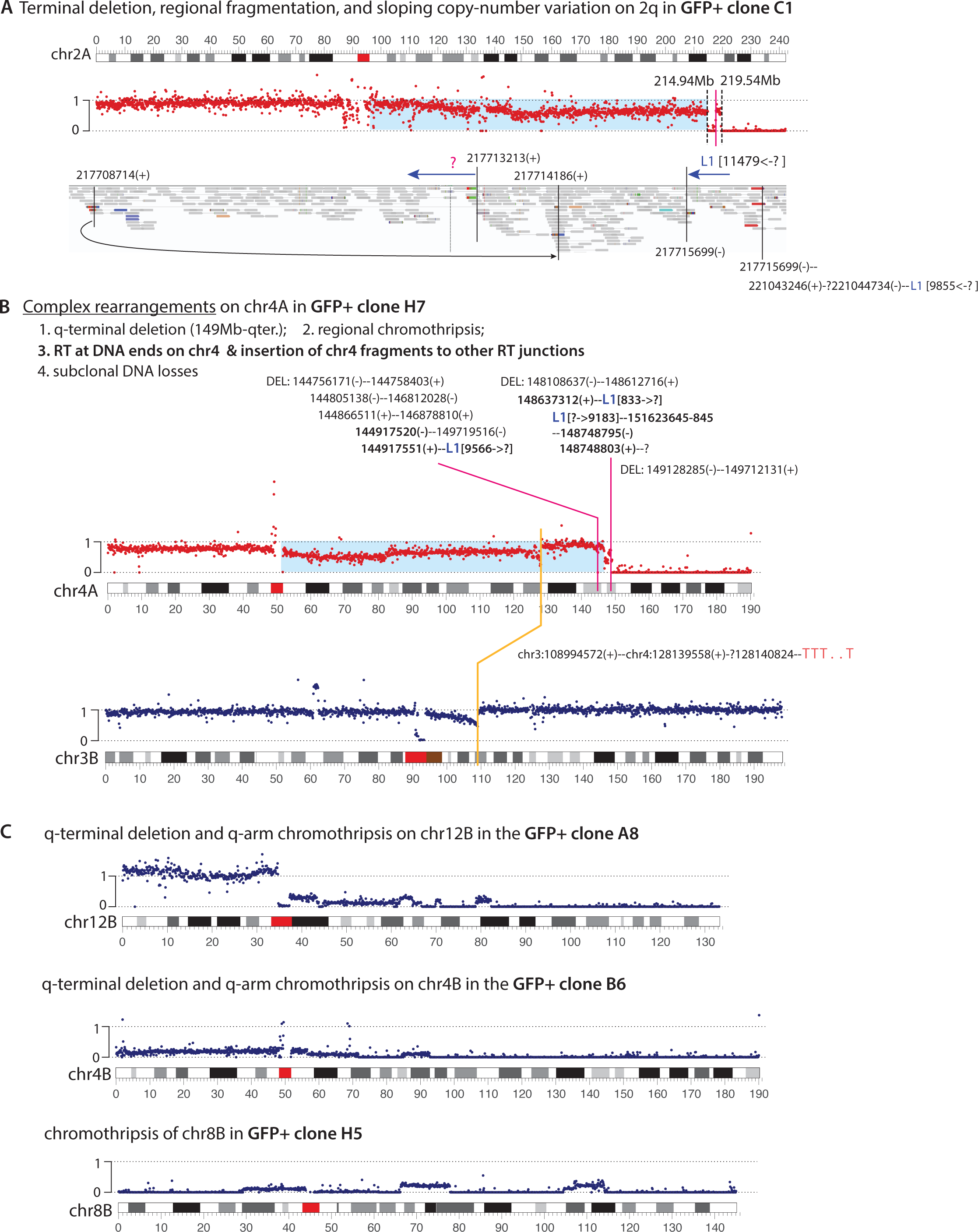
Additional examples of chromothripsis in GFP+ L1 clones. **A.** An example consistent with the outcome of BFB cycles with regional chromosome fragmentation (214.9-219.5Mb). L1 insertions are identified at three junctions (blue arrows showing the orientation of the first cDNA strand) with the following breakpoints: 217,713,213(+) joining a polyadenylated L1, 217,715,699(-) and 221,044,734(-) both joining the 5’-ends of L1 insertions. **B.** A similar example as in **A**. L1 insertions are found at three junctions involving the following breakpoints: 144,917,551(+) and 148,637,312(+) both joining 3’-ends of a non-polyadenylated L1 sequences, 148,748,795(-) joining the 5’-end of an L1 sequence. These breakpoints and their reciprocal breakpoints are highlighted in bold. We also identify a junction involving the breakpoint chr4:128,140,824(-) that joins a poly-T sequence. **C.** Three examples of chromothripsis indicated by oscillating DNA deletion and retention. Breakpoints are unmapped.

## Notes

### Competing Interest Statement

The authors have declared no competing interest.

### Summary of Updates

We added examples of L1-mediated rearrangements from cancer genomes that have been subjected to long-read sequencing. Newly added data are found in Fig. 1B, Figure S3, Figure 4, Fig. 5B, Fig. S4A, Figure 6, Fig. S5D, F, and Fig. S7A.

## References

1. Lander, E.S., Linton, L.M., Birren, B., Nusbaum, C., Zody, M.C., Baldwin, J., Devon, K., Dewar, K., Doyle, M., FitzHugh, W., et al. (2001). Initial sequencing and analysis of the human genome. Nature 409, 860–921. 10.1038/35057062.

2. Burns, K.H., and Boeke, J.D. (2012). Human transposon tectonics. Cell 149, 740–752. 10.1016/j.cell.2012.04.019.

3. Richardson, S.R., Doucet, A.J., Kopera, H.C., Moldovan, J.B., Garcia-Perez, J.L., and Moran, J.V. (2015). The Influence of LINE-1 and SINE Retrotransposons on Mammalian Genomes. Microbiology spectrum 3, MDNA3-0061-2014. 10.1128/microbiolspec.MDNA3-0061-2014.

4. Moran, J.V., Holmes, S.E., Naas, T.P., DeBerardinis, R.J., Boeke, J.D., and Kazazian, H.H., Jr. (1996). High frequency retrotransposition in cultured mammalian cells. Cell 87, 917–927. 10.1016/s0092-8674(00)81998-4.

5. Morrish, T.A., Gilbert, N., Myers, J.S., Vincent, B.J., Stamato, T.D., Taccioli, G.E., Batzer, M.A., and Moran, J.V. (2002). DNA repair mediated by endonuclease-independent LINE-1 retrotransposition. Nature genetics 31, 159–165. 10.1038/ng898.

6. Holmes, S.E., Singer, M.F., and Swergold, G.D. (1992). Studies on p40, the leucine zipper motif-containing protein encoded by the first open reading frame of an active human LINE-1 transposable element. J Biol Chem 267, 19765–19768.

7. Khazina, E., Truffault, V., Buttner, R., Schmidt, S., Coles, M., and Weichenrieder, O. (2011). Trimeric structure and flexibility of the L1ORF1 protein in human L1 retrotransposition. Nature structural & molecular biology 18, 1006–1014. 10.1038/nsmb.2097.

8. Weichenrieder, O., Repanas, K., and Perrakis, A. (2004). Crystal structure of the targeting endonuclease of the human LINE-1 retrotransposon. Structure 12, 975–986. 10.1016/j.str.2004.04.011.

9. Baldwin, E.T., van Eeuwen, T., Hoyos, D., Zalevsky, A., Tchesnokov, E.P., Sánchez, R., Miller, B.D., Di Stefano, L.H., Ruiz, F.X., Hancock, M., et al. (2024). Structures, functions and adaptations of the human LINE-1 ORF2 protein. Nature 626, 194–206. 10.1038/s41586-023-06947-z.

10. Thawani, A., Ariza, A.J.F., Nogales, E., and Collins, K. (2024). Template and target-site recognition by human LINE-1 in retrotransposition. Nature 626, 186–193. 10.1038/s41586-023-06933-5.

11. Luan, D.D., Korman, M.H., Jakubczak, J.L., and Eickbush, T.H. (1993). Reverse transcription of R2Bm RNA is primed by a nick at the chromosomal target site: a mechanism for non-LTR retrotransposition. Cell 72, 595–605. 10.1016/0092-8674(93)90078-5.

12. Ghanim, G.E., Hu, H., Boulanger, J., and Nguyen, T.H.D. (2025). Structural mechanism of LINE-1 target-primed reverse transcription. Science (New York, N.Y.) 388, eads8412. 10.1126/science.ads8412.

13. Jin, W., Yu, C., Zhang, Y., Cao, C., Xia, T., Song, G., Cai, Z., Xue, Y., Zhu, B., and Xu, R.M. (2025). Mechanism of DNA targeting by human LINE-1. Science (New York, N.Y.) 390, eadu3433. 10.1126/science.adu3433.

14. McIntyre, J.J.R., Horton, C.A., and Collins, K. (2025). Different repair pathways support intact or truncated insertions by R2 retrotransposon protein. Science (New York, N.Y.), eadz3121. 10.1126/science.adz3121.

15. Nam, C.H., Youk, J., Kim, J.Y., Lim, J., Park, J.W., Oh, S.A., Lee, H.J., Park, J.W., Won, H., Lee, Y., et al. (2023). Widespread somatic L1 retrotransposition in normal colorectal epithelium. Nature 617, 540–547. 10.1038/s41586-023-06046-z.

16. Burns, K.H. (2017). Transposable elements in cancer. Nature reviews. Cancer 17, 415–424. 10.1038/nrc.2017.35.

17. Mendez-Dorantes, C., and Burns, K.H. (2023). LINE-1 retrotransposition and its deregulation in cancers: implications for therapeutic opportunities. Genes & development 37, 948–967. 10.1101/gad.351051.123.

18. Rodic, N., Sharma, R., Sharma, R., Zampella, J., Dai, L., Taylor, M.S., Hruban, R.H., Iacobuzio-Donahue, C.A., Maitra, A., Torbenson, M.S., et al. (2014). Long interspersed element-1 protein expression is a hallmark of many human cancers. The American journal of pathology 184, 1280–1286. 10.1016/j.ajpath.2014.01.007.

19. Taylor, M.S., Wu, C., Fridy, P.C., Zhang, S.J., Senussi, Y., Wolters, J.C., Cajuso, T., Cheng, W.C., Heaps, J.D., Miller, B.D., et al. (2023). Ultrasensitive Detection of Circulating LINE-1 ORF1p as a Specific Multicancer Biomarker. Cancer discovery 13, 2532–2547. 10.1158/2159-8290.Cd-23-0313.

20. Chalitchagorn, K., Shuangshoti, S., Hourpai, N., Kongruttanachok, N., Tangkijvanich, P., Thong-ngam, D., Voravud, N., Sriuranpong, V., and Mutirangura, A. (2004). Distinctive pattern of LINE-1 methylation level in normal tissues and the association with carcinogenesis. Oncogene 23, 8841–8846. 10.1038/sj.onc.1208137.

21. Doucet-O’Hare, T.T., Rodić, N., Sharma, R., Darbari, I., Abril, G., Choi, J.A., Young Ahn, J., Cheng, Y., Anders, R.A., Burns, K.H., et al. (2015). LINE-1 expression and retrotransposition in Barrett’s esophagus and esophageal carcinoma. Proceedings of the National Academy of Sciences of the United States of America 112, E4894–4900. 10.1073/pnas.1502474112.

22. Lee, E., Iskow, R., Yang, L., Gokcumen, O., Haseley, P., Luquette, L.J., 3rd, Lohr, J.G., Harris, C.C., Ding, L., Wilson, R.K., et al. (2012). Landscape of somatic retrotransposition in human cancers. Science 337, 967–971. 10.1126/science.1222077.

23. Solyom, S., Ewing, A.D., Rahrmann, E.P., Doucet, T., Nelson, H.H., Burns, M.B., Harris, R.S., Sigmon, D.F., Casella, A., Erlanger, B., et al. (2012). Extensive somatic L1 retrotransposition in colorectal tumors. Genome research 22, 2328–2338. 10.1101/gr.145235.112.

24. Helman, E., Lawrence, M.S., Stewart, C., Sougnez, C., Getz, G., and Meyerson, M. (2014). Somatic retrotransposition in human cancer revealed by whole-genome and exome sequencing. Genome research 24, 1053–1063. 10.1101/gr.163659.113.

25. Tubio, J.M.C., Li, Y., Ju, Y.S., Martincorena, I., Cooke, S.L., Tojo, M., Gundem, G., Pipinikas, C.P., Zamora, J., Raine, K., et al. (2014). Mobile DNA in cancer. Extensive transduction of nonrepetitive DNA mediated by L1 retrotransposition in cancer genomes. Science (New York, N.Y.) 345, 1251343. 10.1126/science.1251343.

26. Rodić, N., Steranka, J.P., Makohon-Moore, A., Moyer, A., Shen, P., Sharma, R., Kohutek, Z.A., Huang, C.R., Ahn, D., Mita, P., et al. (2015). Retrotransposon insertions in the clonal evolution of pancreatic ductal adenocarcinoma. Nature medicine 21, 1060–1064. 10.1038/nm.3919.

27. Rodriguez-Martin, B., Alvarez, E.G., Baez-Ortega, A., Zamora, J., Supek, F., Demeulemeester, J., Santamarina, M., Ju, Y.S., Temes, J., Garcia-Souto, D., et al. (2020). Pan-cancer analysis of whole genomes identifies driver rearrangements promoted by LINE-1 retrotransposition. Nature genetics 52, 306–319. 10.1038/s41588-019-0562-0.

28. Gasior, S.L., Wakeman, T.P., Xu, B., and Deininger, P.L. (2006). The human LINE-1 retrotransposon creates DNA double-strand breaks. Journal of molecular biology 357, 1383–1393. 10.1016/j.jmb.2006.01.089.

29. Ardeljan, D., Steranka, J.P., Liu, C., Li, Z., Taylor, M.S., Payer, L.M., Gorbounov, M., Sarnecki, J.S., Deshpande, V., Hruban, R.H., et al. (2020). Cell fitness screens reveal a conflict between LINE-1 retrotransposition and DNA replication. Nature structural & molecular biology 27, 168–178. 10.1038/s41594-020-0372-1.

30. Zumalave, S., Santamarina, M., Espasandín, N.P., Garcia-Souto, D., Temes, J., Baker, T.M., Pequeño-Valtierra, A., Otero, I., Rodríguez-Castro, J., Oitabén, A., et al. (2024). Synchronous L1 retrotransposition events promote chromosomal crossover early in human tumorigenesis. bioRxiv, 2024.2008.2027.596794. 10.1101/2024.08.27.596794.

31. Zhang, C.-Z., Mendez-Dorantes, C., Burns, K.H., and Pellman, D. (2026). A breakage–replication/fusion process explains complex rearrangements and segmental DNA amplification. Nature genetics. 10.1038/s41588-025-02434-5.

32. An, W., Dai, L., Niewiadomska, A.M., Yetil, A., O’Donnell, K.A., Han, J.S., and Boeke, J.D. (2011). Characterization of a synthetic human LINE-1 retrotransposon ORFeus-Hs. Mobile DNA 2, 2. 10.1186/1759-8753-2-2.

33. Rangwala, S.H., and Kazazian, H.H., Jr. (2009). The L1 retrotransposition assay: a retrospective and toolkit. Methods 49, 219–226. 10.1016/j.ymeth.2009.04.012.

34. Umbreit, N.T., Zhang, C.Z., Lynch, L.D., Blaine, L.J., Cheng, A.M., Tourdot, R., Sun, L., Almubarak, H.F., Judge, K., Mitchell, T.J., et al. (2020). Mechanisms generating cancer genome complexity from a single cell division error. Science (New York, N.Y.) 368. 10.1126/science.aba0712.

35. Esnault, C., Maestre, J., and Heidmann, T. (2000). Human LINE retrotransposons generate processed pseudogenes. Nature genetics 24, 363–367. 10.1038/74184.

36. Cooke, S.L., Shlien, A., Marshall, J., Pipinikas, C.P., Martincorena, I., Tubio, J.M., Li, Y., Menzies, A., Mudie, L., Ramakrishna, M., et al. (2014). Processed pseudogenes acquired somatically during cancer development. Nature communications 5, 3644. 10.1038/ncomms4644.

37. Chu, C., Borges-Monroy, R., Viswanadham, V.V., Lee, S., Li, H., Lee, E.A., and Park, P.J. (2021). Comprehensive identification of transposable element insertions using multiple sequencing technologies. Nature communications 12, 3836. 10.1038/s41467-021-24041-8.

38. Miller, T.L.A., Orpinelli Rego, F., Buzzo, J.L.L., and Galante, P.A.F. (2021). sideRETRO: a pipeline for identifying somatic and polymorphic insertions of processed pseudogenes or retrocopies. Bioinformatics 37, 419–421. 10.1093/bioinformatics/btaa689.

39. Bao, C., Tourdot, R.W., Brunette, G.J., Stewart, C., Sun, L., Baba, H., Watanabe, M., Agoston, A.T., Jajoo, K., Davison, J.M., et al. (2023). Genomic signatures of past and present chromosomal instability in Barrett’s esophagus and early esophageal adenocarcinoma. Nature communications 14, 6203. 10.1038/s41467-023-41805-6.

40. Achom, M., Sadagopan, A., Bao, C., McBride, F., Li, J., Konda, P., Tourdot, R.W., Xu, Q., Nakhoul, M., Gallant, D.S., et al. (2024). A genetic basis for sex differences in Xp11 translocation renal cell carcinoma. Cell 187, 5735–5752 e5725. 10.1016/j.cell.2024.07.038.

41. Richardson, C., and Jasin, M. (2000). Frequent chromosomal translocations induced by DNA double-strand breaks. Nature 405, 697–700. 10.1038/35015097.

42. Mendez-Dorantes, C., Kalinowski, J.C., Law, C.T., Schofield, P., Burn, A., and Burns, K.H. (2025). L1 insertion intermediates recombine with one another or with DNA breaks to form genome rearrangements. bioRxiv. 10.1101/2025.09.17.676864.

43. Ostertag, E.M., and Kazazian, H.H., Jr. (2001). Twin priming: a proposed mechanism for the creation of inversions in L1 retrotransposition. Genome research 11, 2059–2065. 10.1101/gr.205701.

44. Biehs, R., Steinlage, M., Barton, O., Juhasz, S., Kunzel, J., Spies, J., Shibata, A., Jeggo, P.A., and Lobrich, M. (2017). DNA Double-Strand Break Resection Occurs during Non-homologous End Joining in G1 but Is Distinct from Resection during Homologous Recombination. Molecular cell 65, 671–684 e675. 10.1016/j.molcel.2016.12.016.

45. Yu, Y., Wang, X., Fox, J., Li, Q., Yu, Y., Hastings, P.J., Chen, K., and Ira, G. (2025). RPA and Rad27 limit templated and inverted insertions at DNA breaks. Nucleic acids research 53. 10.1093/nar/gkae1159.

46. Gilbert, N., Lutz, S., Morrish, T.A., and Moran, J.V. (2005). Multiple fates of L1 retrotransposition intermediates in cultured human cells. Molecular and cellular biology 25, 7780–7795. 10.1128/mcb.25.17.7780-7795.2005.

47. Gilbert, N., Lutz-Prigge, S., and Moran, J.V. (2002). Genomic deletions created upon LINE-1 retrotransposition. Cell 110, 315–325. 10.1016/s0092-8674(02)00828-0.

48. Symer, D.E., Connelly, C., Szak, S.T., Caputo, E.M., Cost, G.J., Parmigiani, G., and Boeke, J.D. (2002). Human l1 retrotransposition is associated with genetic instability in vivo. Cell 110, 327–338. 10.1016/s0092-8674(02)00839-5.

49. Zhou, Y., Caron, P., Legube, G., and Paull, T.T. (2014). Quantitation of DNA double-strand break resection intermediates in human cells. Nucleic acids research 42, e19. 10.1093/nar/gkt1309.

50. Ait Saada, A., Guo, W., Costa, A.B., Yang, J., Wang, J., and Lobachev, K.S. (2023). Widely spaced and divergent inverted repeats become a potent source of chromosomal rearrangements in long single-stranded DNA regions. Nucleic acids research 51, 3722–3734. 10.1093/nar/gkad153.

51. Carvalho, C.M., Ramocki, M.B., Pehlivan, D., Franco, L.M., Gonzaga-Jauregui, C., Fang, P., McCall, A., Pivnick, E.K., Hines-Dowell, S., Seaver, L.H., et al. (2011). Inverted genomic segments and complex triplication rearrangements are mediated by inverted repeats in the human genome. Nature genetics 43, 1074–1081. 10.1038/ng.944.

52. Li, Y., Roberts, N.D., Wala, J.A., Shapira, O., Schumacher, S.E., Kumar, K., Khurana, E., Waszak, S., Korbel, J.O., Haber, J.E., et al. (2020). Patterns of somatic structural variation in human cancer genomes. Nature 578, 112–121. 10.1038/s41586-019-1913-9.

53. Scully, R., Panday, A., Elango, R., and Willis, N.A. (2019). DNA double-strand break repair-pathway choice in somatic mammalian cells. Nature reviews. Molecular cell biology 20, 698–714. 10.1038/s41580-019-0152-0.

54. Ly, P., Brunner, S.F., Shoshani, O., Kim, D.H., Lan, W., Pyntikova, T., Flanagan, A.M., Behjati, S., Page, D.C., Campbell, P.J., and Cleveland, D.W. (2019). Chromosome segregation errors generate a diverse spectrum of simple and complex genomic rearrangements. Nature genetics 51, 705–715. 10.1038/s41588-019-0360-8.

55. Maciejowski, J., Li, Y., Bosco, N., Campbell, P.J., and de Lange, T. (2015). Chromothripsis and Kataegis Induced by Telomere Crisis. Cell 163, 1641–1654. 10.1016/j.cell.2015.11.054.

56. Zhang, C.Z., Spektor, A., Cornils, H., Francis, J.M., Jackson, E.K., Liu, S., Meyerson, M., and Pellman, D. (2015). Chromothripsis from DNA damage in micronuclei. Nature 522, 179–184. 10.1038/nature14493.

57. McClintock, B. (1942). The Fusion of Broken Ends of Chromosomes Following Nuclear Fusion. Proc Natl Acad Sci U S A 28, 458–463. 10.1073/pnas.28.11.458.

58. McClintock, B. (1951). Chromosome organization and genic expression. Cold Spring Harbor symposia on quantitative biology 16, 13–47.

59. McClintock, B. (1950). The origin and behavior of mutable loci in maize. Proc Natl Acad Sci U S A 36, 344–355.

60. Curcio, M.J., and Derbyshire, K.M. (2003). The outs and ins of transposition: from mu to kangaroo. Nat Rev Mol Cell Biol 4, 865–877. 10.1038/nrm1241.

61. Kazazian, H.H., Jr. (2004). Mobile elements: drivers of genome evolution. Science 303, 1626–1632. 10.1126/science.1089670.

62. Carbone, L., Harris, R.A., Gnerre, S., Veeramah, K.R., Lorente-Galdos, B., Huddleston, J., Meyer, T.J., Herrero, J., Roos, C., Aken, B., et al. (2014). Gibbon genome and the fast karyotype evolution of small apes. Nature 513, 195–201. 10.1038/nature13679.

63. Lien, S., Koop, B.F., Sandve, S.R., Miller, J.R., Kent, M.P., Nome, T., Hvidsten, T.R., Leong, J.S., Minkley, D.R., Zimin, A., et al. (2016). The Atlantic salmon genome provides insights into rediploidization. Nature 533, 200–205. 10.1038/nature17164.

64. Ghezraoui, H., Piganeau, M., Renouf, B., Renaud, J.B., Sallmyr, A., Ruis, B., Oh, S., Tomkinson, A.E., Hendrickson, E.A., Giovannangeli, C., et al. (2014). Chromosomal translocations in human cells are generated by canonical nonhomologous end-joining. Molecular cell 55, 829–842. 10.1016/j.molcel.2014.08.002.

65. Youk, J., Kwon, H.W., Lim, J., Kim, E., Kim, T., Kim, R., Park, S., Yi, K., Nam, C.H., Jeon, S., et al. (2024). Quantitative and qualitative mutational impact of ionizing radiation on normal cells. Cell Genom 4, 100499. 10.1016/j.xgen.2024.100499.

66. Zhao, M., Presila, B., Zhang, C.-Z., and Alexander, S. (2025). Ionizing Radiation Shapes Genome Evolution Through Nuclear Abnormalities That Trigger Delayed Proliferative Death. bioRxiv, 2025.2012.2001.691539. 10.64898/2025.12.01.691539.

67. Symington, L.S. (2014). End resection at double-strand breaks: mechanism and regulation. Cold Spring Harbor perspectives in biology 6. 10.1101/cshperspect.a016436.

68. Kloosterman, W.P., Francioli, L.C., Hormozdiari, F., Marschall, T., Hehir-Kwa, J.Y., Abdellaoui, A., Lameijer, E.W., Moed, M.H., Koval, V., Renkens, I., et al. (2015). Characteristics of de novo structural changes in the human genome. Genome research 25, 792–801. 10.1101/gr.185041.114.

69. Al-Zain, A.M., Nester, M.R., Ahmed, I., and Symington, L.S. (2023). Double-strand breaks induce inverted duplication chromosome rearrangements by a DNA polymerase delta-dependent mechanism. Nature communications 14, 7020. 10.1038/s41467-023-42640-5.

70. Ramsden, D.A., Carvajal-Garcia, J., and Gupta, G.P. (2022). Mechanism, cellular functions and cancer roles of polymerase-theta-mediated DNA end joining. Nature reviews. Molecular cell biology 23, 125–140. 10.1038/s41580-021-00405-2.

71. McKerrow, W., Wang, X., Mendez-Dorantes, C., Mita, P., Cao, S., Grivainis, M., Ding, L., LaCava, J., Burns, K.H., Boeke, J.D., and Fenyö, D. (2022). LINE-1 expression in cancer correlates with p53 mutation, copy number alteration, and S phase checkpoint. Proceedings of the National Academy of Sciences of the United States of America 119. 10.1073/pnas.2115999119.

72. Pisanic, T.R., 2nd, Asaka, S., Lin, S.F., Yen, T.T., Sun, H., Bahadirli-Talbott, A., Wang, T.H., Burns, K.H., Wang, T.L., and Shih, I.M. (2019). Long Interspersed Nuclear Element 1 Retrotransposons Become Deregulated during the Development of Ovarian Cancer Precursor Lesions. The American journal of pathology 189, 513–520. 10.1016/j.ajpath.2018.11.005.

73. Xia, Z., Cochrane, D.R., Tessier-Cloutier, B., Leung, S., Karnezis, A.N., Cheng, A.S., Farnell, D.A., Magrill, J., Schmidt, D., Kommoss, S., et al. (2019). Expression of L1 retrotransposon open reading frame protein 1 in gynecologic cancers. Human pathology 92, 39–47. 10.1016/j.humpath.2019.06.001.

74. Sato, S., Gillette, M., de Santiago, P.R., Kuhn, E., Burgess, M., Doucette, K., Feng, Y., Mendez-Dorantes, C., Ippoliti, P.J., Hobday, S., et al. (2023). LINE-1 ORF1p as a candidate biomarker in high grade serous ovarian carcinoma. Sci Rep 13, 1537. 10.1038/s41598-023-28840-5.

